# Deoxysphingolipids activate cGAS-STING1 and enhance antitumor immunity

**DOI:** 10.1101/2024.10.16.618749

**Authors:** Suchandrima Saha, Fabiola N. Velazquez, Fatemeh Dehpanah, Brianna L. Bergfalk, Samantha Mulkeen, Karissa J. Lopez, Cungui Mao, David C. Montrose

**Affiliations:** Department of Pathology, Renaissance School of Medicine, Stony Brook University, Stony Brook, NY; Department of Medicine, Renaissance School of Medicine, Stony Brook University, Stony Brook, NY; Stony Brook Cancer Center, Stony Brook, NY

**Keywords:** colorectal cancer, mitochondria, chemokines, amino acids

## Abstract

Deoxysphingolipids (deoxySLs) are a class of non-canonical sphingolipids that can adversely impact mitochondrial function. Mitochondrial disruption can activate the cyclic GMP–AMP synthase–stimulator of interferon genes (cGAS–STING1) pathway, and in turn, promote antitumor immunity. Here, we investigated whether inducing deoxySL accumulation in colon cancer cells drives immune-specific, antitumor effects. We show that depleting serine increased deoxySLs and enhanced immune cell infiltration in colon tumors, in parallel with suppressing tumor growth. Elevating tumor deoxySL levels through mutation of SPT in cells and feeding a high alanine diet to mice exerted similar effects, including immune-mediated tumor growth suppression. Work conducted in multiple *in vitro* systems identified deoxySLs as key triggers of cGAS-STING1 activation, an effect mediated by mitochondrial release of double-stranded DNA resulting from mitochondrial stress. Collectively, these findings provide the first evidence that deoxySLs can drive antitumor immunity and identify a previously unrecognized metabolic strategy for cancer therapy.

## Introduction

Canonical sphingolipids (SLs) are a class of bioactive lipids generated through condensation of the non-essential amino acid serine and palmitoyl-CoA by serine-palmitoyl transferase (SPT) ^1–3^. Although originally believed to simply be structural components of cell membranes and organelles, numerous studies have revealed their additional role as signaling molecules ^2,4^. Under low serine conditions, an increased ratio of alanine to serine or mutation of SPT (causing it to favor alanine over serine as a substrate), non-canonical SLs, known as deoxySLs, are produced ^5,6^. Intracellular accumulation of deoxySLs can become toxic, as evidenced by neuropathies driven by inherent mutations in SPT which adversely impact peripheral neurons ^7,8^. Recent work suggests that deoxySL-driven mitochondrial toxicity may be responsible for these adverse effects ^9,10^. In addition to these neurologic and mitochondrial-related findings, recent studies have highlighted their potent anticancer properties, including suppression of anchorage-independent cancer cell proliferation, cancer spheroid growth, and tumor growth in an immunodeficient setting ^11,12^. However, it is unknown whether inducing deoxySL accumulation impacts tumor immunity.

The cyclic GMP-AMP Synthase-Stimulator of Interferon Genes (cGAS-STING) pathway is capable of driving robust antitumor immunity, through its induction of a signal transduction cascade leading to phosphorylation of the transcription factor interferon regulatory factor 3 (IRF3) and subsequent Type I interferon (IFN) secretion ^13–15^. cGAS-STING activation is triggered by the presence of cytosolic double-stranded DNA (dsDNA), which in addition to originating from pathogens, can be derived from mitochondria ^16,17^. Mitochondrial DNA (mtDNA) release into the cytosol occurs as a result of mitochondrial membrane permeabilization, often driven by mitochondrial stress ^16,18,19^. In fact, the ability of mtDNA release to trigger cGAS-STING has been highlighted by multiple studies showing that mtDNA depletion prevents Type I IFN production during mitochondrial disruption ^20–22^. The level of mitochondrial stress capable of inducing mtDNA release can be triggered by altered intracellular metabolism resulting from nutrient deprivation or direct disruption of mitochondrial membranes ^20,23,24^. However, the optimal approach to trigger mitochondrial stress in cancer cells to induce cGAS-STING driven antitumor immunity remains to be established.

Given that deoxySLs can induce mitochondrial disruption, the current study investigates whether they stimulate antitumor immunity through the cGAS-STING1 pathway. Here, we demonstrate that increasing deoxySL abundance in colon cancer cells activates cGAS-STING1 and triggers downstream production of Type I IFNs through mitochondrial dysfunction and subsequent mtDNA release into the cytosol. We further show that increasing intratumoral deoxySLs through serine deprivation or SPT mutation of tumor cells combined with a high alanine diet drives immune-mediated tumor growth suppression in mice. Taken together, these findings provide the first evidence of an immune-specific role of deoxySLs for conferring their anticancer effects.

## Results

### Serine depletion elevates tumor deoxySLs and increases antitumor immunity in colorectal cancer models

Our previous work demonstrated that exogenous and endogenous deprivation of serine drives immune-mediated tumor growth suppression ^20^. Given that serine depletion induces accumulation of non-canonical deoxySLs ^11,12,25,26^, we posited that deoxySLs may be causally linked to these effects. To test this possibility, we first measured the levels of total deoxydihydroCeramides (deoxydhCeramides) in subcutaneous CT26 tumors with shRNA-mediated knockdown of Luciferase (*shLuc*) from mice fed complete (Comp) diet (serine sufficient conditions) and tumors with knockdown of *Psat1* (*shPsat1*), a key serine synthesis enzyme, from mice fed serine deficient (Ser Def) diet (serine deficient conditions), as previously described^20^. Here, we found significantly increased levels of total deoxydhCeramides in serine deplete tumors, largely driven by very long chain species, while no changes in canonical SLs were observed (Figures 1A-C). We next measured SL levels in serine sufficient *vs*. deficient conditions in an autochthonous colon tumor model, established through azoxymethane injections of mice carrying shRNAs targeting *Renilla* or *Psat1*, as previously described ^27^. Here, we found that reducing serine availability to colon tumors through *Psat1* knockdown (Figure S1) and feeding a Ser Def diet for 3 weeks increased total deoxydhCeramide levels approximately 8-fold (Figure 1D). No changes in canonical SLs were observed (Figures 1E and F). In association with the observed deoxySL-specific increase, *Ifnb1* expression and abundance of CD4^+^ T helper and CD8^+^ cytotoxic T cells were significantly elevated in tumors, which corresponded to suppression of growth (Figures 1G-I). Collectively, these findings suggest a role for deoxySLs in eliciting antitumor immunity upon serine deprivation.

**Figure 1.**
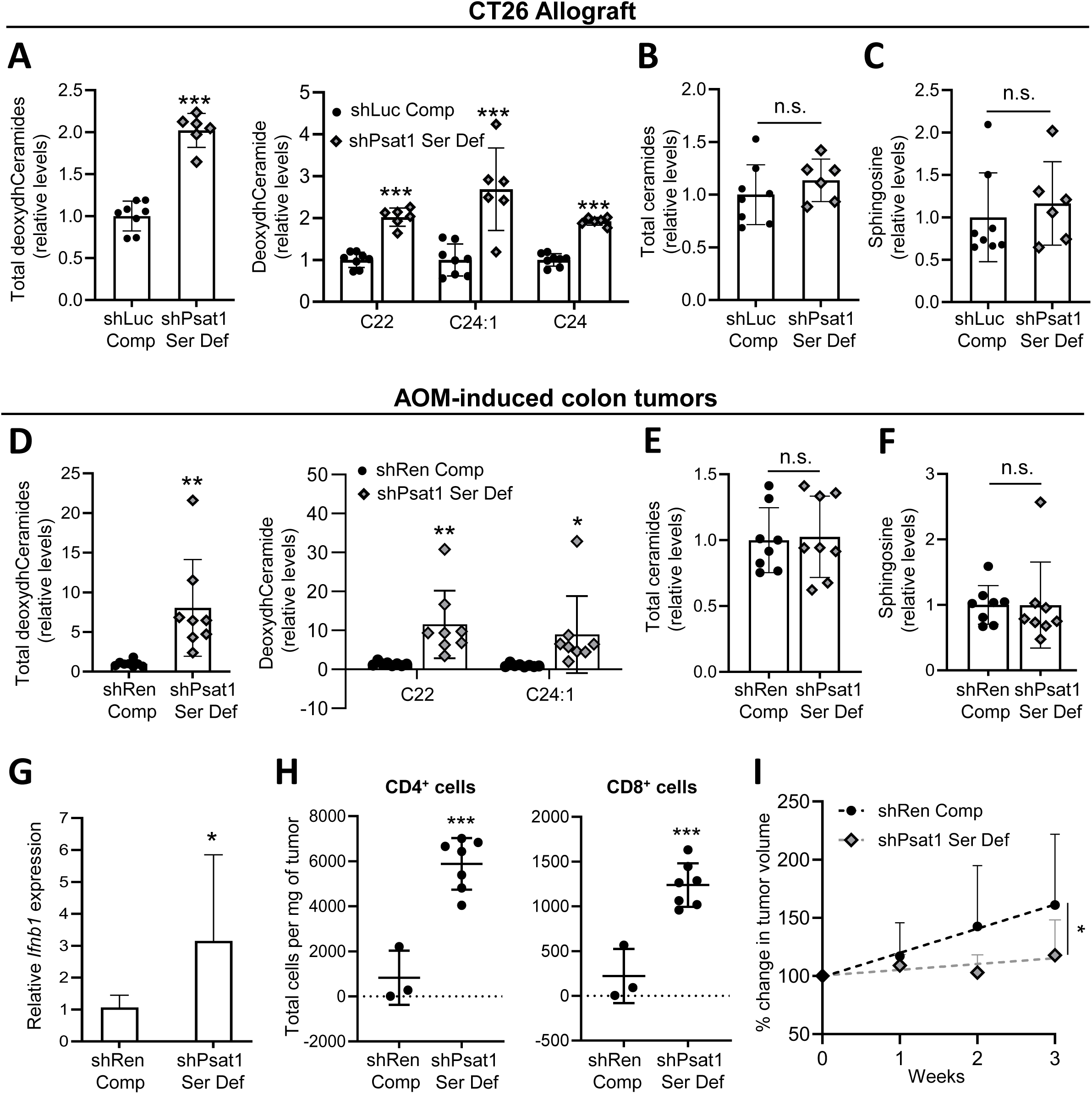
Serine depletion increases deoxysphingolipids and immune infiltration in colon tumors. A-C, CT26 cells carrying doxycycline (doxy)-inducible shRNAs targeting *Luciferase* (*shLuc*) or *Psat1* (*shPsat1*) were injected subcutaneously into separate groups of BALB/c mice and allowed to form a tumor over 14 days while mice were fed amino acid-based complete (Comp) diet. Mice bearing *shPsat1* tumors were then switched to a serine deficient (Ser Def) diet while *shLuc* tumor-bearing mice were continued on Comp diet. Doxy was given by oral gavage every other day to all mice to activate shRNAs beginning on day 14. Abundance of total deoxydihydroCeramides (deoxydhCeramides) (left side) and individual deoxydhCeramide species (right side) (A), total ceramides (B), and sphingosine (C) was measured in tumors on day 18 (n=5 mice/group) and displayed as relative to the *shLuc* Comp group. D-I, Mice carrying shRNAs targeting *Renilla* (*shRen*) or *Psat1* (*shPsat1*) were administered 6 weekly injections of azoxymethane while being fed AIN93G diet. After visible colon tumor formation (six weeks after the last injection), *shRen* mice were fed Comp diet and *shPsat1* mice were fed Ser Def diet and all mice were orally gavaged with doxy every other day for the following 3 weeks. Tumors were harvested and the abundance of total deoxydhCeramides (left side) and individual deoxydhCeramide species (right side) (D), total ceramides (E), and sphingosine (F) was measured and displayed as relative to the *shRen* Comp group, gene expression of *Ifnb1* was determined (G), the number of CD4^+^ T and CD8^+^ T was quantified (H), and the change in tumor volume was calculated relative to Week 0, based on images captured by colonoscopy (I) (n=10-12 mice/group). Data represent mean values ± S.D. *P≤0.05; **P≤0.01; ***P≤0.001 (paired t-tests).

### DeoxySLs mediate cGAS-STING1 activation following serine depletion

Given the data shown above and recent evidence that serine deprivation activates the immune-stimulating cGAS-STING1 pathway ^20^, we next tested whether its induction can be driven by deoxySL production in the context of serine depletion. First, the levels of total deoxydhCeramides were measured in serine-depleted CT26 cells *in vitro*, which showed deoxydhCeramide accumulation compared to serine sufficient conditions, primarily driven by very long chain species (Figures 2A and B). Notably, serine deprivation did not elevate the concentrations of the canonical sphingolipids sphingosine and total ceramide (Figure S2). To mechanistically connect deoxySL generation with the activation of the cGAS-STING1 pathway, cells were simultaneously deprived of serine and treated with the SPT inhibitor myriocin (Myr) to block deoxySL production. Myr treatment strongly attenuated deoxySL accumulation upon serine deprivation in parallel with blocking increases in the cGAS-STING1 components, phosphorylated TBK1, STING and IRF3, and expression of cGAS-STING1-induced genes (Figures 2C-F; Figure S3). Taken together, these findings suggest that cGAS-STING1 pathway activation following serine depletion is mediated by deoxySL accumulation.

**Figure 2.**
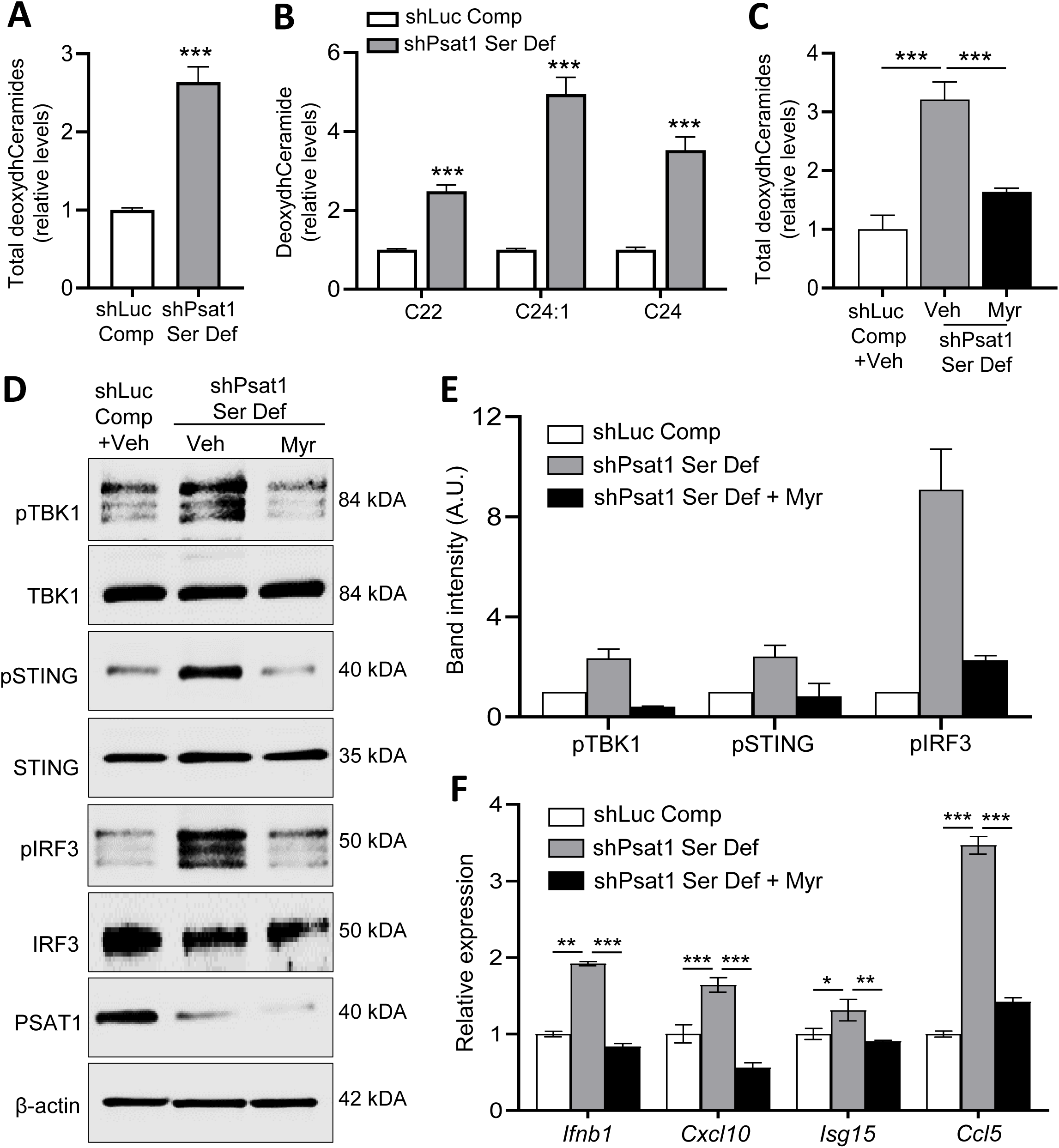
Deoxysphingolipids activate the cGAS-STING1 pathway following serine depletion in colon cancer cells. A-B, CT26 cells carrying doxycycline (doxy)-inducible shRNAs targeting Luciferase (*shLuc*) or Psat1 (*shPsat1)* were cultured in complete (Comp) or serine deficient (Ser Def) medium and treated with doxy for 24 h and the abundance of total deoxydihydroCeramides (deoxydhCeramides) (A) and individual deoxydhCeramide species (B) were measured in cells and displayed as relative to the *shLuc* Comp group. C-F, *shLuc* or *shPsat1* cells were cultured in Comp or Ser Def medium, respectively, and treated with vehicle (Veh) (0.1% methanol) or 500 nM myriocin (Myr), as indicated, for 24 h and the abundance of total deoxydhCeramides was determined and displayed as relative to the *shLuc* Comp + Veh group (C), western blotting was carried out to probe for the indicated proteins (D), the fold change in expression of phosphorylated proteins shown in panel d was determined from two independent experiments (E), and the expression of the indicated genes was measured (F). Data represent mean values ± S.D. *P≤0.05; **P≤0.01; ***P≤0.001 (paired t-tests).

### Deoxysphinganine treatment activates the cGAS-STING1 pathway to induce a Type I IFN response

To more directly examine whether deoxySLs are capable of inducing cGAS-STING1 activation, WT CT26 cells cultured in standard medium were treated with 1-deoxysphinganine (dSA), the precursor for deoxydhceramide biosynthesis, which selectively drives deoxySL production in cells (Figure S4) ^25^. Here, we found that dSA increased the protein expression of phosphorylated TBK1, STING and IRF3 and gene expression of *Ifnb1*, *Cxcl10*, *Isg15* and *Ccl5*, as well as the secretion of IFNβ1 (Figures 3A-D; Figure S5). Similar observations were made in dSA-treated DLD1 human colon cancer cells (Figure S6). To next confirm whether the cGAS-STING pathway is responsible for the dSA-mediated increase in *Ifnb1* and downstream response genes, *Sting1* was knocked down in dSA-treated cells (Figure 3E). Here, blocking STING1 activation completely abrogated the dSA-mediated increase in downstream Type I IFN responsive gene expression (Figure 3F). Altogether, these findings demonstrate that deoxySLs mediate cGAS-STING1 activation to drive the Type I IFN response.

**Figure 3.**
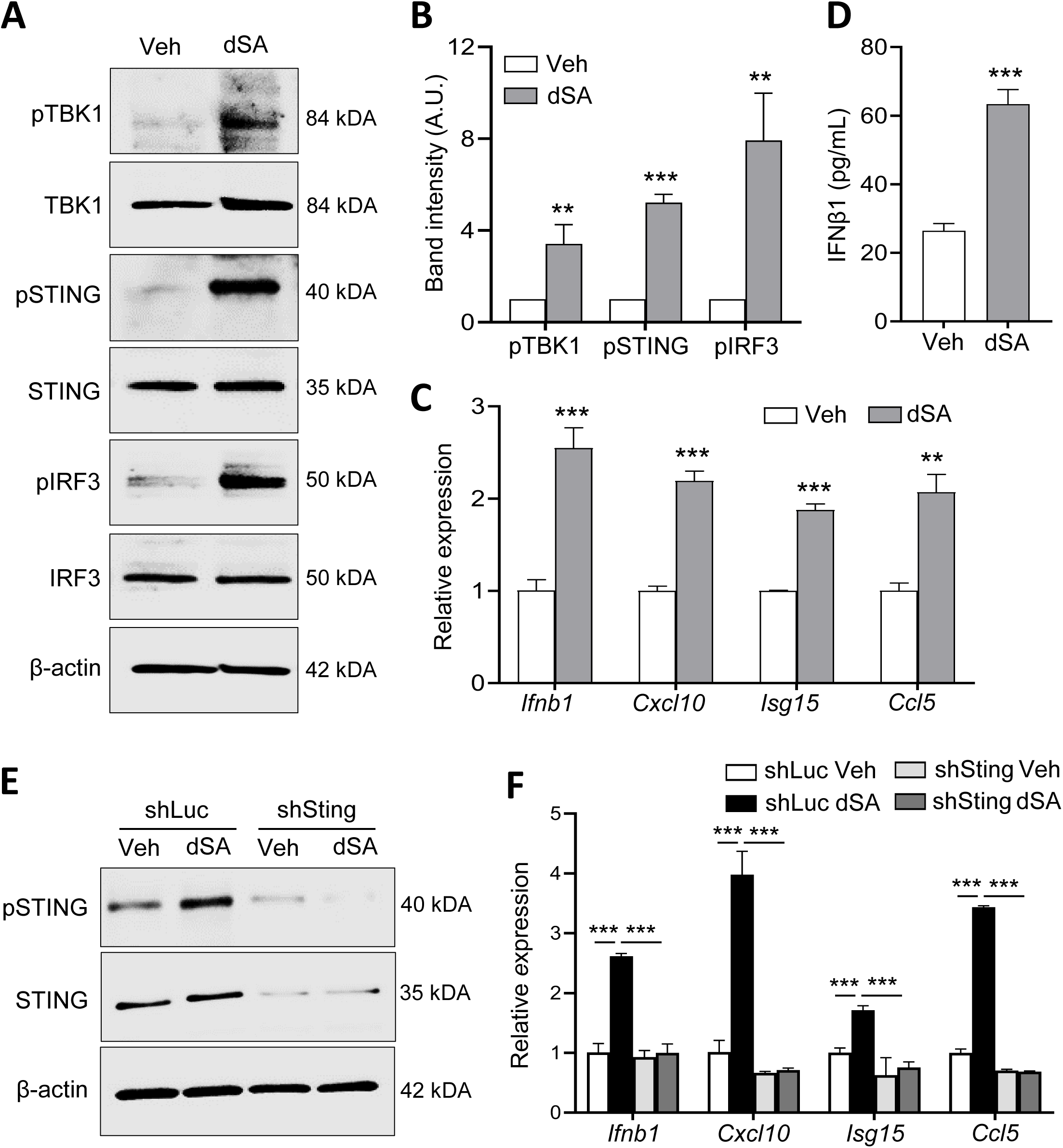
Deoxysphinganine treatment of colon cancer cells induces the Type I IFN response through cGAS-STING1. A-C, WT CT26 cells were treated with vehicle (Veh) (DMSO) or 500 nM 1-deoxysphinganine (dSA) for 18 h and western blotting was carried out to probe for the indicated proteins (A), the fold change in expression of phosphorylated proteins was determined from 3 independent experiments (B), the expression of the indicated genes was determined (C) and the levels of IFNβ1 were measured in medium (D). E-F, CT26 cells carrying doxycycline (doxy)-inducible shRNAs targeting *Luciferase* (*shLuc*) or *Sting1* (*shSting*) were given doxy for 24 h, followed by treatment with Veh or 500 nM dSA for 18 h and the expression of the indicated proteins (E) and the indicated genes (F), were measured. Data represent mean values ± S.D. *P≤0.05; **P≤0.01; ***P≤0.001 (paired t-tests).

### Deoxysphingolipids activate cGAS-STING1 by inducing cytosolic accumulation of mitochondrial DNA

Mitochondrial dysfunction leads to release of its DNA into the cytosol, which can activate cGAS-STING1 ^16,17^. Therefore, we first examined whether dSA treatment induces mitochondrial dysfunction in colon cancer cells by measuring mitochondrial membrane potential (MMP) and reactive oxygen species (ROS). Analysis revealed significantly reduced MMP and increased ROS in CT26 cells after exposure to 500 nM dSA for 6 hours (Figures 4A and B). Similar effects were observed in DLD1 cells (Figure S7). Despite these mitochondrial defects, this dose of dSA did not trigger apoptosis up to 24 hours after treatment (Figure S8). In parallel with the observed dSA-induced mitochondrial dysfunction, cytosolic dsDNA accumulated, which colocalized with the mitochondrial markers COXIV and TFAM, while less than 10% associated with the nucleus, suggesting its mitochondrial origin (Figures 4C-E; Figure S9). To directly demonstrate that deoxySL-mediated induction of mtDNA release is responsible for cGAS-STING1 pathway activation, expression of phosphorylated-STING1, TBK1 and IRF3 and downstream Type I interferon responsive genes were measured in rho^0^ CT26 cells (depleted of mtDNA). Here, treatment with dSA failed to activate cGAS-STING1 components or increase the expression of *Ifnb1* and associated response genes (Figures 4F-H; Figure S10). Altogether, these findings support the concept that deoxySL-induced mtDNA release and cytosolic accumulation in colon cancer cells lead to the activation of the innate immune responsive cGAS-STING1 pathway.

**Figure 4.**
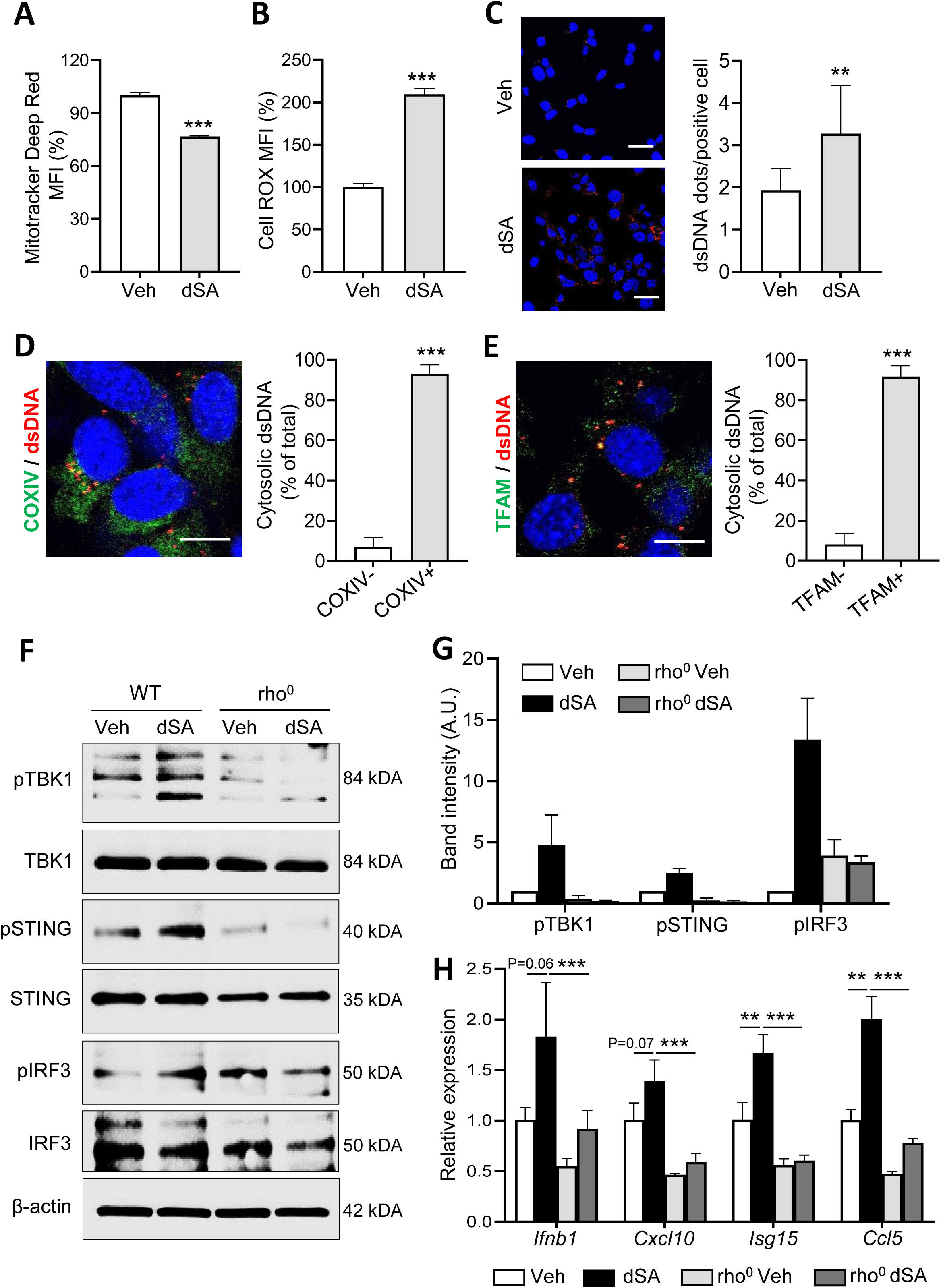
Deoxysphinganine triggers release of mitochondrial DNA, leading to activation of cGAS-STING1 in colon cancer cells. A-B, WT CT26 cells were treated with 500 nM 1-deoxysphinganine (dSA) or vehicle (Veh) (DMSO) for 6 h and mitochondrial membrane potential was determined by flow cytometry after staining with Mitotracker Deep Red (A) and reactive oxygen species levels were determined by flow cytometry after staining with CellROX (B). Values in the Veh group were set to 100% and values for the dSA-treated group are displayed as relative to Veh. C-E, Cells were treated as described in panel a and cytosolic dsDNA accumulation was assessed by immunofluorescence using a dsDNA-specific antibody (red color). Representative images (left side) and quantification of the percent of cells that were positive for cytosolic dsDNA (right), are shown (C). Cells were co-stained for dsDNA and COXIV and representative images (left side) and quantification of the percent of cytosolic dsDNA dots in proximity to COXIV (right side), are shown (D). Cells were stained for dsDNA and TFAM and representative images (left side) and quantification of the percent of cytosolic dsDNA co-localizing with TFAM, are shown (E). DAPI (blue color) was used to stain the nuclei in all images. Scale bar = 10 μm. F-H, CT26 WT and rho^0^ cells were treated with dSA of Veh for 18 h and extracted protein was subjected to western blotting to probe for the indicated proteins (F), the fold change in expression of phosphorylated proteins shown in panel f was determined from two independent experiments (G), and the relative expression of the indicated genes was determined (H). Data represent mean values ± S.D. **P≤0.01; ***P≤0.001 (paired t-tests).

### SPT mutation and alanine supplementation activate cGAS-STING1

SPT mutation (which drives utilization of alanine over serine as a substrate for making SLs) or alanine supplementation to WT cells elevates deoxySLs ^11,28,29^. Therefore, we next examined the impact of these approaches on cGAS-STING1 activation, alone or in combination. First, we showed that wild-type *SPTLC1* (SPT WT) DLD1 cells supplemented with alanine or mutant *SPTLC1* (SPT MUT) cells cultured in control medium exhibited increased deoxySLs, while the combination enhanced this effect (Figure 5A (left side); Figure S11A). The same conditions resulted in similar effects in CT26 cells (Fig. 5A (right side); Figure S11B). We next determined whether deoxySL accumulation in this setting was associated with induction of cGAS-STING1. Modest increases in phosphorylated TBK1, STING and IRF3 and expression of IRF3 target genes *Ifnb1*, *Cxcl10*, and *Isg15* were observed with alanine supplementation in WT cells and SPT mutation alone, while these effects were markedly enhanced in alanine-supplemented SPT MUT cells (Figures 5B and C). Collectively, these findings further establish a role for deoxySLs in activating cGAS-STING1 signaling in colon cancer cells.

**Figure 5.**
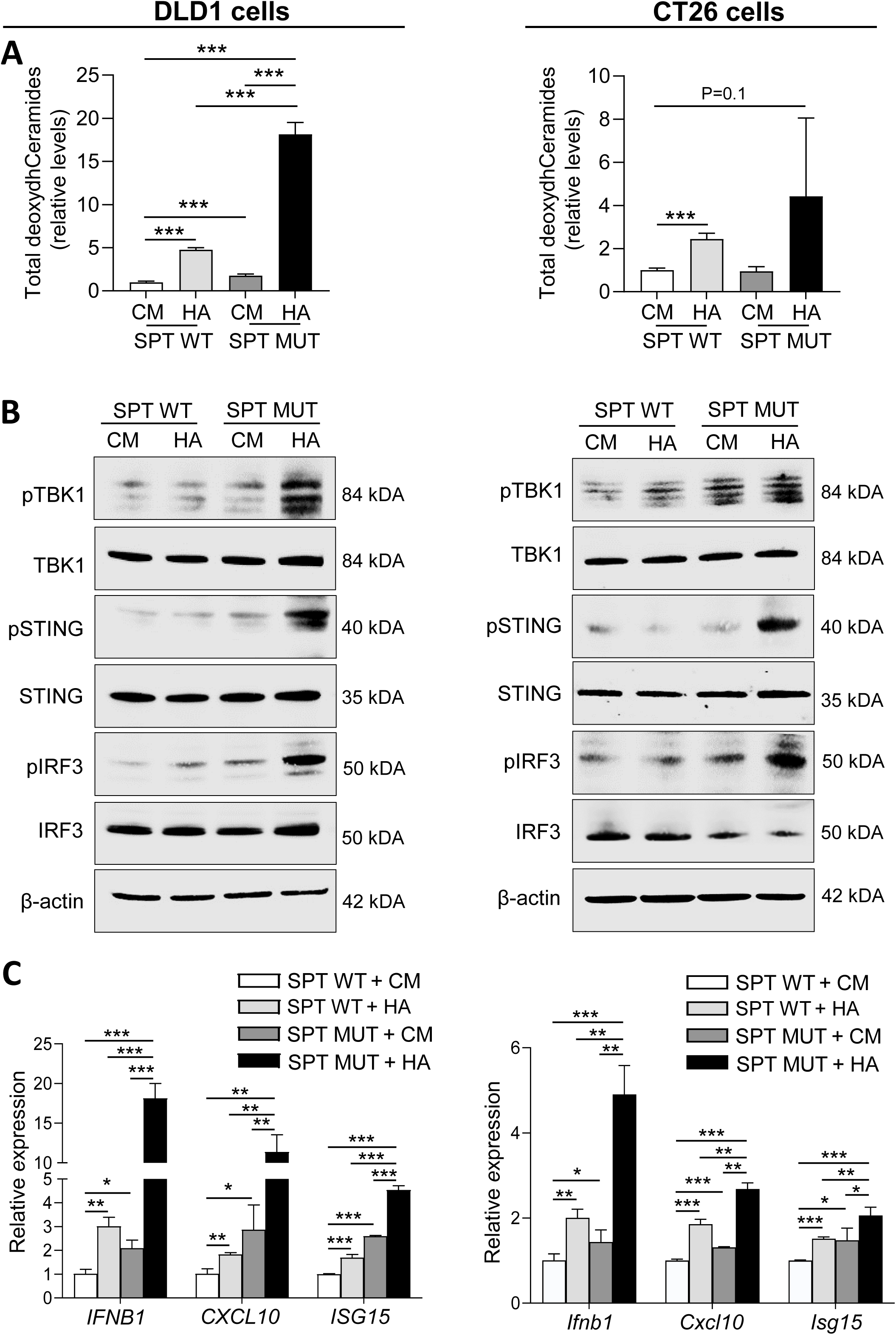
Serine palmitoyl transferase mutation and alanine supplementation promote the activation of cGAS-STING1 pathway and Type I interferon response. A, Doxycycline (doxy)-inducible *SPTLC1* wild-type (SPT WT) and *SPTLC1^C133W^* mutant (SPT MUT) CT26 and DLD1 cells were treated with doxy for 24 h then cultured in complete medium (CM) or high alanine medium (HA) as indicated, and the abundance of total deoxydihydroCeramides (deoxydhCeramides) was quantified and displayed as relative to SPT WT cells given CM, after 30 h in DLD1 cells (left side) and 14 h in CT26 cells (right side) (n = 4 samples per group). B, CT26 and DLD1 cells were treated as described in panel a and the levels of the indicated proteins were determined for DLD1 (left side) and CT26 cells (right side). C, CT26 and DLD1 cells were treated as described in panel a and the expression of the indicated genes were measured after 36 h in DLD1 (left side) and 24 h in CT26 cells (right side). Data represent mean values ± S.D. *P≤0.05; **P≤0.01; ***P≤0.001 (paired t-tests).

### Inducing deoxySL production triggers immune-mediated tumor suppression

We next investigated whether the observed activation of cGAS-STING1 induced by SPT mutation and alanine supplementation in colon cancer cells translates to immune stimulation *in vivo*. To test this possibility, SPT WT and MUT CT26 cells were inoculated into the flanks of separate groups of syngeneic BALB/c mice, and after palpable tumors were established, mice were equally randomized to receive a control amino acid-based diet (CD) or high alanine diet (HAD) (Table S1) and administered doxycycline (doxy) to activate WT or mutant SPT (Figure 6A). SPT MUT tumors from mice fed CD and SPT WT tumors from mice fed HAD exhibited increased levels of deoxydhCeramides compared to SPT WT tumors from mice given CD, while combining SPT mutation with a HAD maximally elevated deoxySLs compared to either condition alone (Figures 6B-E). In contrast, no changes in canonical sphingolipids were observed (Figures 6F and G). Tumor volume measurements revealed that SPT MUT tumors from mice fed CD and SPT WT tumors from mice fed HAD had suppressed growth, while this effect was enhanced when SPT mutation was combined with the HAD, compared to control conditions (Figure 7A). Tumor weight measurements at the end of the study confirmed these observations (Figure 7B). We next determined whether the observed tumor suppression and deoxySL accumulation were associated with alterations in immune cell populations within tumors. Flow cytometry-based measurements revealed that SPT mutant tumors from mice fed HAD (which exhibited the strongest tumor suppression and highest deoxydhCeramide levels) showed an approximate doubling of activated dendritic cells (DCs) and activated CD8^+^ T cells compared to SPT WT tumors from mice fed CD (Figure 7C). A trend for increased activated DCs were observed in SPT WT tumors from mice fed HAD and SPT MUT tumors from mice fed CD, although these differences were not significant (Figure 7C). To determine whether the observed increase in tumor immune infiltration when combining SPT mutation and HAD is causally linked to tumor growth suppression, mice bearing SPT WT tumors and fed CD or SPT MUT tumors and fed HAD were treated with either control IgG or CD4^+^ and CD8^+^ depleting antibodies. Here, we found that systemic depletion of CD4^+^ and CD8^+^ T cells completely abrogated the anti-neoplastic effects of SPT mutation and dietary alanine supplementation (Figures 7D and E; Figure S12). These results further substantiate that deoxySL accumulation suppresses the growth of tumors by eliciting an antitumor immune response.

**Figure 6.**
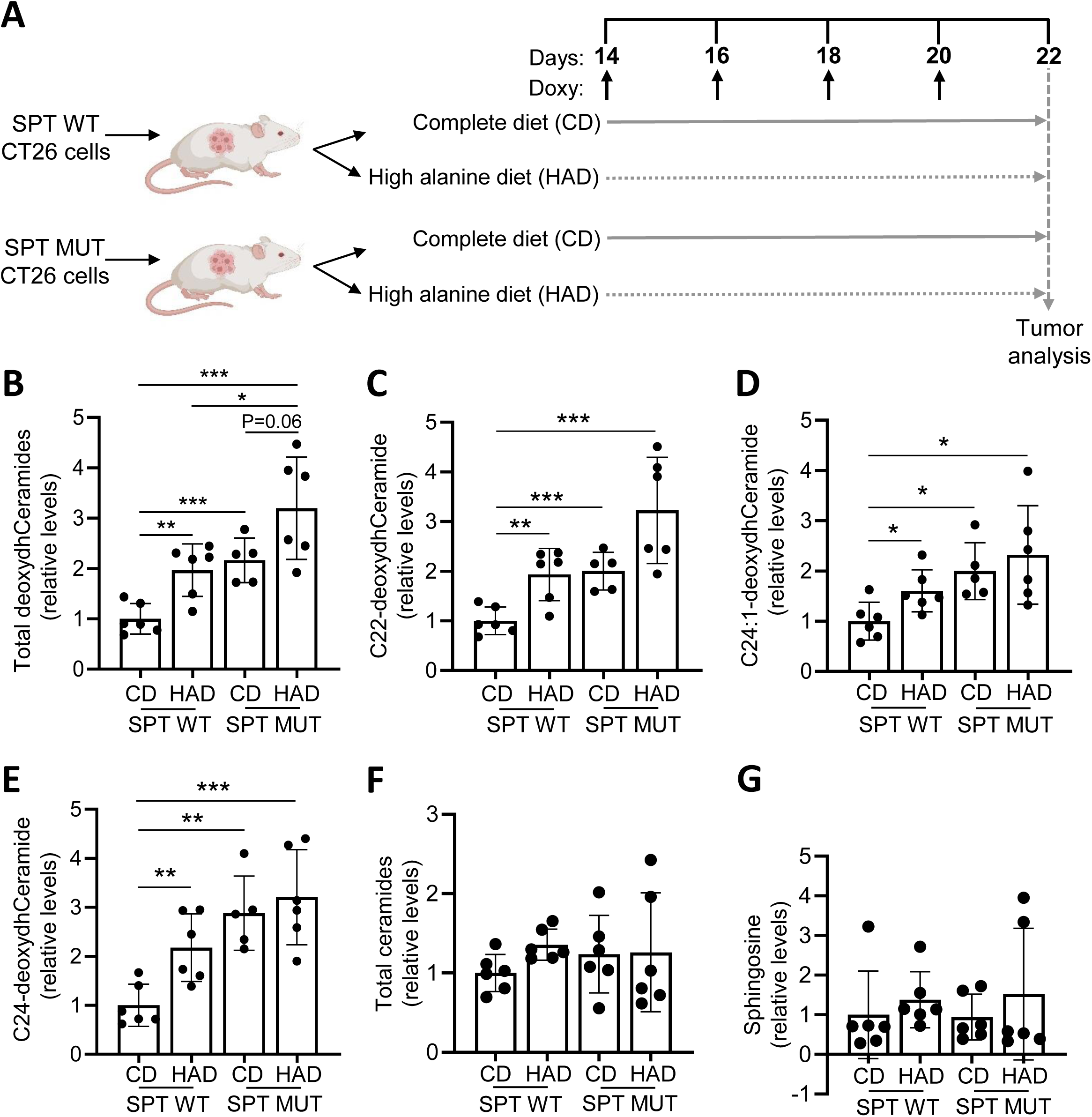
Serine palmitoyl transferase mutation and high alanine feeding promotes deoxysphingolipid production in tumors. A, Schematic depicting experimental design whereby CT26 cells carrying doxycycline (doxy)-inducible wild-type *SPTLC1* (SPT WT) or mutant *SPTLC1^C133W^* (SPT MUT) were inoculated into the flanks of BALB/c mice and tumors were allowed to develop over 2 weeks. Half of the mice bearing either tumor type were fed control amino acid based diet (CD) and the other half were fed high alanine diet (HAD), while all mice received doxy by oral gavage every other day. Tumors were harvested in Day 22 and utilized for analysis. B-G, The abundance of total deoxydihydroceramides (deoxydhceramides) (B), C22-deoxydhCeramide (C) C24:1-deoxydhCeramide (D), C24-deoxydhCeramide (E), total ceramides (F) and sphingosine (G) were determined and displayed as relative to SPT WT tumors from mice given CD (n=5-6 per group). Data represent mean values ± S.D. *P≤0.05; **P≤0.01; ***P≤0.001 (paired t-tests).

**Figure 7.**
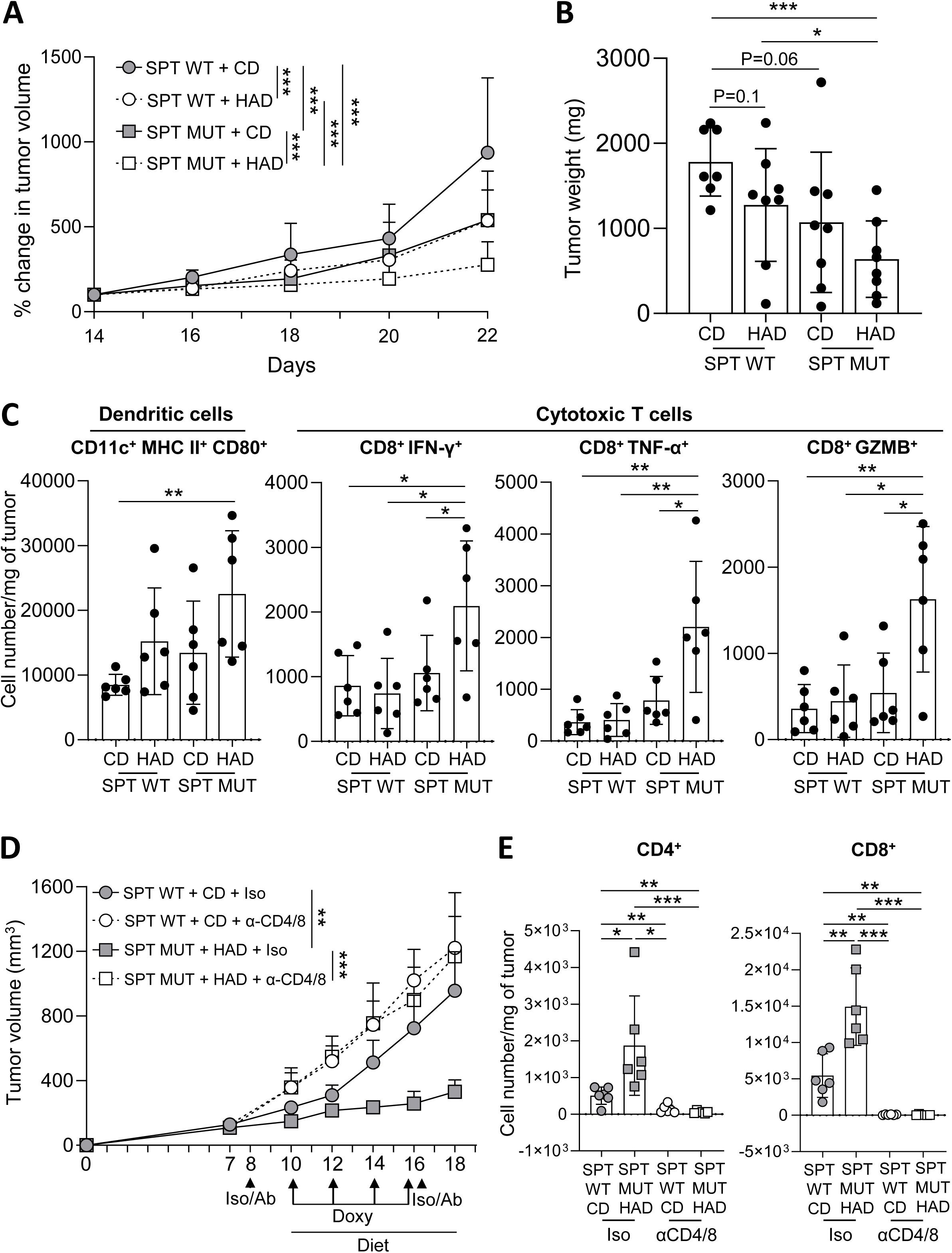
Increasing deoxysphingolipids suppresses tumor growth by enhancing immune cell influx. A-C, Doxycycline (doxy)-inducible *SPTLC1* wild-type (SPT WT) or *SPTLC1^C133W^* mutant (SPT MUT) CT26 cells were injected subcutaneously into separate groups of BALB/c mice and tumors were allowed to develop over 14 days while mice were fed complete diet (CD). Half of the tumor-bearing mice were then randomized to either continue receiving CD or switched to a high alanine diet (HAD), and doxy (20 mg/kg) was given by oral gavage every other day to all mice to activate shRNAs (n=7-8 mice/group). Tumor size was measured every other day and calculated as percent change compared to day 14 (A), tumor weights were measured on day 22 (B), flow cytometry was performed on single cell preparations of tumors on day 22 to measure the number of activated dendritic cells and cytotoxic T cells (C). D-E, SPT WT or SPT MUT CT26 cells were injected subcutaneously into BALB/c mice and tumors were allowed to develop over 14 days while mice were fed CD. Mice bearing WT tumors were continued on CD while mice carrying MUT tumors were switched to HAD, and mice in either group were randomized to receive isotype control (Iso) or CD4^+^ and CD8^+^ blocking antibodies (α-CD4/8) on days 8 and 16, as denoted by arrowheads. Doxy was given by oral gavage every other day to all mice to activate shRNAs, as indicated by arrows. Tumor volume was measured over 18-days (n = 6 mice/group) (D), and flow cytometry was carried out on Day 18 to determine the total number of CD4+ T cells and CD8+ T cells in tumors (n = 6 mice per group) (E). Data represent mean values ± S.D. *P≤0.05; **P≤0.01; ***P≤0.001.

## Discussion

Here, we show that deoxySLs induce mitochondrial dysfunction in colon cancer cells, leading to release of mtDNA and subsequent activation of the cGAS-STING1 pathway. Notably, genetic and diet-mediated approaches to increase deoxySLs in tumors suppress tumor growth in an immune-dependent manner. Taken together, these findings suggest a novel role for non-canonical sphingolipids in driving antitumor immunity, providing potential additional treatment options for colorectal cancer and other cancers.

Numerous studies have shown that depleting exogenous and/or endogenous serine exerts anticancer effects, including improving response to chemotherapy ^30–37^. Recent work extending upon these findings has shown that serine depletion triggers activation of the cGAS-STING1 pathway and accompanying induction of antitumor immunity ^20^. The current work reveals a marked increase in deoxySL levels in *Psat1* deficient tumors from mice fed Ser Def diet compared to control tumors in both autochthonous and allograft models. Notably, this deoxySL elevation was accompanied by significant increases in immune cell influx into tumors and tumor growth suppression, suggesting a link between deoxySLs and antitumor immunity. The fact that treatment of serine-deprived colon cancer cells with myriocin, which brought deoxydhCeramide levels close to what is found under control conditions, blocked cGAS-STING1 induction and the accompanying chemokine expression increase, supports a direct role for deoxySLs in inducing the cGAS-STING1 pathway. Our additional findings that other methods to drive deoxySL accumulation (e.g., dSA treatment, SPT mutation, excess alanine) similarly induced cGAS-STING1 further support a role for this class of SLs in triggering this pathway. Moreover, feeding a HAD to mice bearing SPT MUT tumors enhanced antitumor immunity, providing a connection between deoxySL-mediated induction of cGAS-STING1 and elevated immune infiltration. Notably, this *in vivo* approach reveals a novel strategy for inducing deoxySL accumulation in tumors.

Previous work has shown that deoxySLs exert anticancer effects independent of immune involvement. For example, direct administration of the synthetic dSA analogue enigmol through diet supplementation suppressed the growth of subcutaneous tumors in immunodeficient hosts^38,39^. Additionally, spisulosine, a dSA analogue isolated from the marine mollusc *Mactromeris polynyma*, directly induced apoptosis of a broad range of human cancer cell lines *in vitro* ^40,41^. More recent work demonstrated that accumulation of deoxySLs in cancer cells inhibits anchorage independent growth of tumor cells *in vitro* and restrains tumor growth *in vivo*, all findings made under immunodeficient conditions ^11,12^. In light of these studies, we cannot exclude the possibility that deoxySLs are mediating tumor suppression through direct toxic effects on the tumor cells themselves. However, we showed that the dose of dSA necessary to trigger cGAS-STING1 activation in colon cancer cells in the current study, did not impact cell viability, making it less likely that direct effects on cancer cells are playing a role in suppressing tumor growth. Most importantly, immune cell depletion abrogated the antitumor effects of deoxySLs, which supports immune activation as a key mechanism by which tumor growth is suppressed in this context.

In light of our findings suggesting that increasing deoxySLs activates the immunostimulatory pathway cGAS-STING1 as a result of mitochondrial dysfunction, a remaining question is the mechanism by which mitochondria become dysfunctional in this context. Previous work has found that exogenous administration of labeled dSA results in the accumulation of deoxydhceramide in both the mitochondria and ER and that very-long chain deoxydhCeramides exert cytotoxicity by disrupting mitochondrial integrity ^9,10^. Therefore, it is possible that deoxySL accumulation in our system directly impacts mitochondrial function and/or induces ER stress, with the latter effect resulting in mitochondrial abnormalities. Separate studies have shown that ER stress and the resulting disruptions in calcium homeostasis can elevate mitochondrial calcium levels leading to mitochondrial dysfunction, thus supporting the possibility that the loss of mitochondrial function is an ER-initiated event ^42,43^. In fact, recent work has shown that the synthesis of very-long chain deoxyceramides is crucial for the neurotoxic effect exerted by deoxySLs through the induction of ER stress ^44^. Certainly, more in-depth studies of both mitochondrial and ER biology in the context of deoxySL accumulation in cancer cells are warranted to elucidate these important concepts.

It is well understood that a hereditary SPT mutation in humans results in neurotoxicity stemming from accumulation of deoxySLs in sensory neurons ^7,8^. Additional preclinical work demonstrates that long-term feeding of a serine deficient or alanine-enriched diet results in neuropathy, resulting from neuronal accumulation of deoxySLs ^45,46^. As such, although elevating deoxySLs in tumor cells exerts promising anticancer effects, likely through immune and non-immune mechanisms, concerns remain regarding the potential for associated toxicities. The work conducted in the current study reveals that short-term induction of deoxySLs exerts rapid tumor suppression and accompanying immune infiltration. And although formal analysis of neurological function was not performed, activity and gross motor function of mice in the experimental groups appeared to be equivalent to control mice. Given that longer-term deoxySL accumulation is responsible for the previously reported neuropathies, it is likely that the short-term induction used here would not induce deficits in neurologic function. Therefore, it is possible that implementing this approach clinically on a short-term basis would be feasible, although the optimal approach for increasing tumor deoxSLs would first need to be determined. Impressively, spisulosine was administered intravenously to patients with advanced cancer, and although these studies were phase I trials with the major goal of determining safety and pharmacokinetics, they highlight the ability to directly administer these lipids to patients ^47,48^. To the best of our knowledge, the current study is the first to demonstrate that feeding an alanine-enriched diet increases deoxySLs in tumors and exerts tumor-suppressive effects. Given the potential ease of supplementing a patient’s diet with amino acids, increasing intake of alanine may be a clinically feasible approach to drive deoxySL accumulation in tumors thereby serving as an adjuvant to current therapies.

## Materials and Methods

### Cell culture

Murine colorectal carcinoma CT26 cells were cultured in RPMI medium (Gibco), human embryonic kidney 293T cells (ATCC) and human colorectal adenocarcinoma DLD1 cells were cultured in DMEM (Gibco). Cell lines were authenticated using the short tandem repeat profiling (ATCC). All cell lines were routinely tested for Mycoplasma using the MycoProbe Mycoplasma Detection Kit (R&D Biosystems) and all experiments were performed within 10 passages after thawing. The cells were maintained in medium containing 10% fetal bovine serum (FBS, Gibco) and 100 U mL−1 penicillin sodium and 100 μg mL−1 streptomycin sulfate (Gibco) at 37 °C under 5% CO_2_. Complete (Comp) medium was prepared using MEM (Gibco) containing 10% dialyzed FBS (Gibco), 100 U mL−1 penicillin sodium and 100 μg mL−1 streptomycin sulfate, MEM vitamin mix, 4.5 g L-1 glucose, 0.4 mmol L-1 glycine and 0.4 mmol L-1 serine. Serine-deficient (Ser Def) medium was made with all components except for serine. High alanine (HA) medium mimicked Comp medium but also contained 3.2 mM of alanine.

### Cloning and lentiviral particle production

shRNAs targeting luciferase (*shLuc*) or murine *Psat1* (*shPsat1*) (clone #640) were cloned into a doxycycline (doxy)-inducible EZ-tet-pLKO-Puro vector (Addgene #85966). Lentiviral particles were generated by transfecting 293T cells with 1.5 µg ps-PAX2 (Addgene #12260), 0.5 µg pCMV-VSV-G (Addgene # 8454) plasmids along with 2 µg plasmid specifically targeting the genes of interest using Lipofectamine 2000. 48 and 72 h post transfection, viral supernatants were collected, and the target cells were transduced with viral supernatants containing polybrene (5 µg mL^−1^). One day following infection, medium was changed and the transduced cells were selected with 5 µg mL^−1^ of puromycin for 48 h.

To generate cells carrying inducible shRNAs targeting *Sting1*, shRNAs targeting luciferase (*shLuc*) or murine *Sting1* (clone #20) were cloned into the EZ-tet-pLKO-Hygro vector (Addgene #85972) and lentiviral particles were generated, as previously described ^20^. Viral supernatants were then added to CT26 cells with polybrene (5 µg mL-1) and one day later, medium was changed and cells were treated with 100 µg mL-1 hygromycin for 3 days to select for transduced cells.

To generate inducible *SPTLC1^WT^* and *SPTLC1^C133W^*-expressing CT26 and DLD1 cell lines, plasmids described in ^11^ were used. HEK293T cells were transfected with 250 ng of pCMV-VSVG, 750 ng of pCMV-dR8.2 and 1 μg of pCW57.1_SPTLC1_WT (Addgene Plasmid #181920) or pCW57.1_SPTLC1_C133W (Addgene Plasmid #181921) using Lipofectamine 2000. Viral supernatants were collected 48 h post transfection and CT26 and DLD1 cells were transduced with the viral supernatants containing polybrene (8 µg mL^−1^). One day later, medium was changed, and cells were treated with 2 µg mL^−1^ puromycin for 72h to select for transduced cells. All the engineered inducible cells were cultured in respective media containing 10% tetracycline-free FBS (Genesee Scientific) and 100 U mL^−1^ penicillin sodium and 100 µg mL^−1^ streptomycin sulfate.

### *In vivo* studies

All experiments involving mice were conducted in accordance with the Institutional Animal Care and Use Committee of Stony Brook University (protocol #: 1345543). Six-week-old female BALB/c mice were purchased from The Jackson Laboratory (Bar Harbor, ME). Mice were anaesthetized using isofluorane and injected subcutaneously in the right flank with 1 × 10^6^ *shLuc* or *shPsat1*#640 CT26 cells suspended in a 1:1 mixture of serum-free RPMI medium and matrigel. Mice were fed AIN-93G until tumors reached 200-250 mm^3^, then *shLuc* tumor-bearing mice were placed on complete (Comp) amino acid-based diet (mimicking AIN-93G) and *shPsat1* tumor-bearing mice were given serine deficient (Ser Def) diet ^33^ (Research Diets). Mice were also administered 20 mg Kg^−1^ doxy by oral gavage every other day to induce shRNAs over the 8-day period, and tumor volume was measured using calipers.

For experiments providing dietary alanine supplementation, six-week-old female BALB/c mice were purchased from The Jackson Laboratory (Bar Harbor, ME). Mice were anaesthetized using isofluorane and subcutaneously injected in the right flank with 1 × 10^6^ cells of *SPTLC1^WT^* or *SPTLC1^C133W^* CT26 cells suspended in a 1:1 ratio of serum-free RPMI medium and matrigel. Mice were maintained in AIN-93G diet until tumors reached ∼200 mm^3^. Fourteen days post inoculation, mice were randomized to either receive Comp diet or HAD (Research Diets) (Table S1) for 8 days. All mice were administered 20 mg Kg^−1^ doxy by oral gavage every other day to induce shRNAs over the 8-day period. Tumor volume was measured manually using calipers.

For experiments involving antibodies blocking CD4^+^ and CD8^+^ T cells, each tumor-bearing mouse received 400 µg of anti-CD4 antibody (BioXCell, clone: GK1.5) and 200 µg of anti-CD8 antibody (BioXCell, clone 2.43) or 600 µg of Rat IgG2b (BioXCell, clone: LTF2) by i.p. injection on days 8 and 16 days after cell inoculation.

To induce autochthonous colon tumors, *shRen* and sh*Psat1* mice (generated as previously described ^27^ backcrossed 8 generation to an FVB/NJ background, were injected i.p. with 10 mg Kg^−1^ of azoxymethane (AOM) (Sigma) dissolved in 0.9% NaCl once per week for 6 weeks, while being fed AIN-93G diet. Six weeks following the last AOM injection, *shRen* mice were switched to Comp diet and *shPsat1* mice were given Ser Def diet for 3 weeks, while all mice were administered 20 mg Kg-1 doxycycline (doxy) by oral gavage every other day, to induce the shRNAs during the 3-week period. Colonoscopies were performed weekly over the treatment period of 3 weeks using a 1.9-mm rigid bore endoscope (Karl Storz, Germany). Briefly, mice were anesthetized using isoflurane, colons were flushed using PBS and the endoscope was inserted into the anus of the mouse until the colonic flexure was visible, and videos were recorded while exiting the colon. To measure tumor size, still images of each tumor were generated from videos and were used to measure the area of the tumor in a blinded manner using Image J (version 1.54g). Tumor size changes for each week were calculated relative to week 0. At the end of the treatment period, mice were euthanized and colon tumors were harvested and processed for lipidomics, gene expression measurements or flow cytometry, as described below.

### Analysis of mitochondrial membrane potential (MMP)

DLD1, CT26 WT and CT26 *shLuc* or *shPsat1* cells were seeded at a density of 3 × 10^5^ in 12-well plates. CT26 WT and DLD1 cells were treated with 500 nM of dSA for 6 h, while CT26 *shLuc* cells were treated with Comp media and s*hPsat1* cells were given Ser Def media in the presence or absence of 500 nM myriocin for 24 h. Cells were harvested and incubated with 10 nM MitoTracker™ Deep Red (Invitrogen) for 15 min at 37°C. The cells were washed twice with PBS and counterstained with 1 μg mL-1 4’,6-diamidino-2-phenylindole (DAPI; Sigma-Aldrich) to distinguish between live and dead cells. All the samples were acquired on an LSRII flow cytometer (BD) and analyzed using BD FACSDiva 9.0 software.

### ROS measurements

CT26 WT and DLD1 cells were seeded at a density of 3 × 10^5^ in 12-well plates and treated with 500 nM of dSA for 6 h respectively. Cells were harvested, and incubated with CellROX green (Invitrogen) at a final concentration of 5 μM in serum-free DMEM for 30 min at 37°C. Cells were then washed twice with PBS, resuspended in cold PBS, counterstained with 1 μg mL-1 DAPI to distinguish live and dead cells. The samples were acquired on LSRII flow cytometer (BD) and analyzed using BD FACSDiva 9.0 software.

### Cell viability assay

Wild-type CT26 cells (3 × 10^5^) were seeded on 12-well plates and treated with vehicle (DMSO) or 0.1, 0.5, 1, 2 or 5 μM of dSA for 24 h. Cells were harvested and stained with 20 nmol/L of 3, 30-Dihexyloxacarbocyanine iodide (DiOC6) (Sigma-Aldrich) for 30 minutes at 37°C. Cells were then stained with 1 μg/mL of 40,6-diamidino-2-phenylindole (DAPI) (Sigma-Aldrich) for 5 minutes. Samples were then subjected to flow cytometric analysis using a BD LSRFortressa II flow cytometer with 405 nm (violet) and 488 nm (blue) lasers. DIOC6 positive and DAPI negative cells were considered live, while DIOC6 negative and DAPI positive or only DAPI positive cells or DIOC6 negative cells were considered dead. Data were analyzed using BD FACSDiva 9.0 software.

### Immunofluorescence microscopy

Cells were grown on glass coverslips and following treatment and incubation, were washed twice with PBS and then fixed with 4% paraformaldehyde (Santa Cruz Biotechnology) for 15 mins. Cells were then permeabilized with 0.1% Tween20 and 0.01% Triton X-100 in PBS on ice, blocked in 2% BSA for 1 h, and incubated with primary antibodies specific for dsDNA (Santa Cruz, 1:200 dilution), LMNB (ab16048, Abcam, 1:500 dilution), COXIV (ab16056, Abcam, 1:300 dilution) or TFAM (GTX103231, GeneTex, 1:500 dilution). Next, cells were washed three times, 5 mins each and incubated with anti-mouse secondary antibody conjugated to Alexa Fluor 594 (ab150120, Invitrogen, 1:500 dilution), or anti-rabbit secondary antibody conjugated to Alexa Fluor 488 (A-11008, Thermo Fisher Scientific, 1:500 dilution) for 2 h at room temperature. Cells were washed three times with PBS, 5 mins each and counterstained using 1 μg mL-1 DAPI to identify the nucleus. Coverslips were then mounted onto slides using Fluoromount G Anti-fade (Southern Biotech) and images were captured using a Leica TCS SP8 X confocal microscope at 100X magnification. Cytosolic dsDNA was quantified using NIH ImageJ (version 1.53k), as previously described ^20^.

### mtDNA depletion

CT26 WT cells were cultured in RPMI medium supplemented with tetracycline-free FBS, 500 ng mL^−1^ ethidium bromide (Sigma-Aldrich), 50 µg mL^−1^ uridine and 1 mM sodium pyruvate for 14 days ^20^, and subjected either to vehicle (DMSO) or 500 nM of dSA for 24 h and harvested for western blots and quantitative real-time PCR.

### Immunoblotting

Cells were harvested and lysed using RIPA buffer (Sigma-Aldrich) supplemented with 1 each of protease and phosphatase inhibitor tablet per 10 mL (ThermoFisher Scientific). The samples were incubated on ice for 15 mins, sonicated for 10 mins and centrifuged at 15,000g for 15 min at 4 °C. The supernatants were then collected and protein concentration was determined using the DC Protein Assay Reagent (Biorad). Equal amounts of proteins were resolved on 4-20% Criterion TGX Precast Midi protein gradient gels (Biorad), transferred to nitrocellulose membranes, blocked with 5% BSA for 1 h at room temperature and incubated overnight at 4°C with primary antibodies targeting phospho-TBK1 (Cell Signaling, 5483, 1:1,000 dilution), TBK1 (Cell Signaling, 3504, 1:2,000 dilution), phopsho-STING (Cell Signaling, 19781, 1:2,000 dilution), STING (Cell Signaling, 13647, 1:2000 dilution) phospho-IRF3 (Cell Signaling, 29047, 1:2,000 dilution), IRF3 (Cell Signaling, 4302, 1:2,000 dilution), PSAT1 (NBP1-32920, Novus Biologicals, 1:2,000 dilution), or anti-beta-Actin peroxidase (Sigma-Aldrich, A3854, 1:20,000 dilution). Following incubation with primary antibody, the membranes were incubated with horseradish peroxidase-conjugated anti-rabbit (Sigma, A0545, 1:3000 dilution) secondary antibody. Bands were visualized using Clarity Max Western ECL Substrate (Bio-Rad) on a C600 Gel Doc and Western Imaging System operated by cSeries capture v.1.6.8.1110 (Azure Biosystems). Uncropped images of immunoblots are shown in Figures S13 and S14.

### Quantitative real-time PCR (qRT-PCR)

Total RNA from cells was isolated using QIAzol (Qiagen) and the quantity of RNA was determined using a NanoDrop ND-1000 spectrophotometer. 4000 ng of total RNA was used to prepare cDNA using oligo (dT) primers and reverse transcriptase (Quanta). Quantitative real-time PCR was performed using PowerTrack™ SYBR Green Master Mix (ThermoFisher) and gene-specific primers for mouse *Ifnb1* (fwd: 5’-AAGATCAACCTCACCTACAG-3’; rev: 5’-AAAGGCAGTGTAACTCTTCT-3’), Interferon-stimulated gene 15 (*Isg15*) (fwd: 5’-ACAGTGATGCTAGTGGTACA-3’; rev: 5’-AAGACCTCATAGATGTTGCT-3’), C-X-C motif chemokine ligand 10 (*Cxcl10*) (fwd: 5’-AAGTTTACCTGAGCTCTTTT-3’; rev: 5’-AGTATCTTGATAACCCCTTG-3’), and Chemokine (C-C motif) ligand 5 (*Ccl5*) (fwd: 5’-AGTGGGTTCAAGAATACATC-3’; rev: 5’-CTAGGACTAGAGCAAGCAAT-3’). Mouse Beta actin (*ACTB*) was used as a housekeeping gene (fwd: 5’-GGCTGTATTCCCCTCCATCG-3’; rev: 5’-CCAGTTGGTAACAATGCCATGT-3’). Genes were amplified using the Applied Biosystems QuantStudio 7 Flex RT-PCR system and the ΔΔCt method was used to calculate the relative expression.

### IFN beta 1 measurements

CT26 cells were seeded on a 60 mm dish and treated with vehicle (DMSO) or 500 nM of dSA for 24h. Conditioned media were collected after 24 hrs and IFNβ1 levels were measured using a VeriKine mouse IFN Beta ELISA Kit (PBL Assay Science), as per the manufacturer’s recommendation. Absorbance was determined at 450 nm using Molecular Device Spectra Max M5 instrument and concentration was calculated relative to a standard curve and normalized to protein concentration.

### Lipidomics

Cells were washed with cold PBS then scraped into 2 mL of cell extraction mix (2:3 of 70% Isopropanol: Ethanol) and stored at −80°C until lipid extraction. Tumor samples stored at −80°C were transferred to Lysing Matrix A 2 mL tube containing garnet matrix and one ceramic sphere (MP Biomedicals) and minced with dissecting scissors in 500 μL of cell extraction mixture (2:3 of 70% Isopropanol: Ethanol). Samples were then homogenized using the FastPrep-24 lysis system (MP Biomedicals), centrifuged and supernatants were obtained. Cell extraction mixture was added, and samples were stored at −80°C until lipid extraction, as previously described ^49^. For sphingolipid quantification, extracted samples from cells or tissues were spiked with 50 pmol of a corresponding internal standard and extracts were analyzed by the Stony Brook Cancer Center Biological Mass Spectrometry Shared Resource using a Thermo Scientific Quantiva coupled to a vanquish HPLC or Agilent 6490 coupled to an Infinity 1290, as described previously ^50,51^. Lipid levels were normalized to total inorganic phosphate.

### Immune profiling

Tumors were excised, weighed, minced into 1-2 mm^3^ pieces and then placed into a digestion solution containing 0.5 mg mL^−1^ collagenase D (Worthington Biochemical Corporation), 0.01 mg mL^−1^ DNase I (Sigma-Aldrich), and 0.5 mg mL^−1^-dispase (Thermo Fisher Scientific) in RPMI medium for 30 min while shaking at 220 rpm at 37°C. Samples were vortexed and strained through a 70 μm cell strainer. The flow-throughs were collected and centrifuged at 1000 g for 5 min at 4°C. The pellets were resuspended in PBS and ACK lysis buffer (Invitrogen) was added to lyse the red blood cells. Samples were washed with FACS buffer (PBS + 2% FBS) followed by blocking of non-specific binding using FACS buffer for 30 min at room temperature. Live/dead staining was performed using LIVE/DEAD Fixable Aqua Dead Cell Stain (Invitrogen) at 1:500 dilution, and 1:200 dilution of fluorochrome-conjugated antibodies for specific cell surface markers were used as follows: CD45 (Biolegend, clone: 30-F11), CD3ε (Biolegend, clone: 145-2C11), CD8 (Biolegend, clone: 53-6.7), CD25 (Biolegend, clone: PC61), MHC class II (Biolegend, clone: M5/114.15.2), CD11c (Biolegend, clone: N418), and CD80 (Biolegend, clone: 16-10A1) for 30 mins at room temperature in dark. Next, the samples were washed in FACS buffer, fixed using 4% PFA in PBS for 20 mins, permeabilized with 0.5% Tween-20, blocked using 1% FBS diluted in permeabilization buffer for 15 mins and then stained with IFN-γ (Biolegend, clone: XMG1.2), TNF-α (Biolegend, clone: MP6-XT22) and Granzyme B (Biolegend, clone: QA16A02) at 1:200 dilution for 30 mins at room temperature in the dark. The samples were washed using FACS buffers and run on an LSRII flow cytometer (BD) and data were analyzed using the Kalluza C software (version1.1). Gating strategies used to determine activated dendritic and T cell populations are shown in Figure S15.

### Statistical analysis

Differences in tumor growth were determined using an exponential regression model of tumor growth rate over time. The statistical comparisons in other assays were carried out using two-tailed Student’s t-tests. Graphs were created using GraphPad Prism 10 and data were plotted as mean +/− standard deviation. Statistical significance was set at P < 0.05, and all experiments were carried out with at least three biological replicates unless otherwise noted in the figure legend.

## Acknowledgments

We wish to acknowledge the Stony Brook Cancer Center Biological Mass Spectrometry Shared Resource for expert assistance with lipidomics analysis. We thank Drs. Yusuf Hannun, Chiara Luberto, Christopher Clarke and Daniel Canals for numerous helpful discussions.

## Notes

**Grant Support:** This work was supported in part by the Pilot Award Program for Team Science from Stony Brook Cancer Center (D.C.M. and C.M.) and the US National Cancer Institute (NCI) grant R01CA273263 (C.M.).

**Declaration of interests:** The authors declare no potential conflicts of interest.

### Competing Interest Statement

The authors have declared no competing interest.

### Summary of Updates

The manuscript has been revised to elucidate the role of dietary and genetically driven deoxySL accumulation in mediating immune-dependent tumor growth suppression in mice. These findings are highly novel and are supported by multiple innovative approaches to elevate deoxySL levels in tumor cells both in vitro and in vivo, including dietary modulation strategies. The figures have been revised, and the manuscript now includes additional figures to further support the findings. The author list has been updated. Supplemental files have been updated.

