## Supplementary File for "Deoxysphingolipids activate cGAS-STING1 and enhance antitumor immunity"

### SUPPLEMENTARY FIGURES

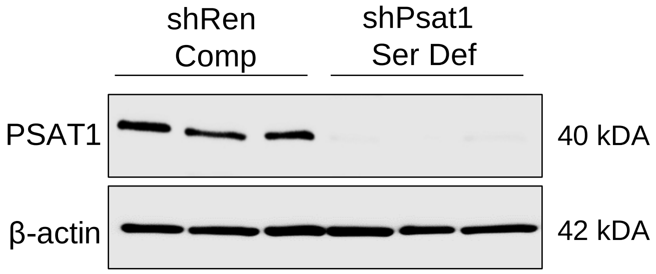


**Figure S1. Doxycycline administration reduces PSAT1 expression in azoxymethane-induced colon tumors from *shPsat1* mice (related to Figure 1)**

Mice carrying shRNAs targeting Renilla (*shRen*) or Psat1 (*shPsat1*) were administered azoxymethane for 6 weeks and after the last injection, tumors were allowed to develop for the next 6 weeks, while mice were fed AIN93G diet. *shRen* mice were then given Complete (Comp) diet and *shPsat1* mice were given Serine Deficient (Ser Def) diet and all mice were orally gavaged with doxy every other day for the next 3 weeks. Tumors were harvested from colons and probed for the indicated proteins.

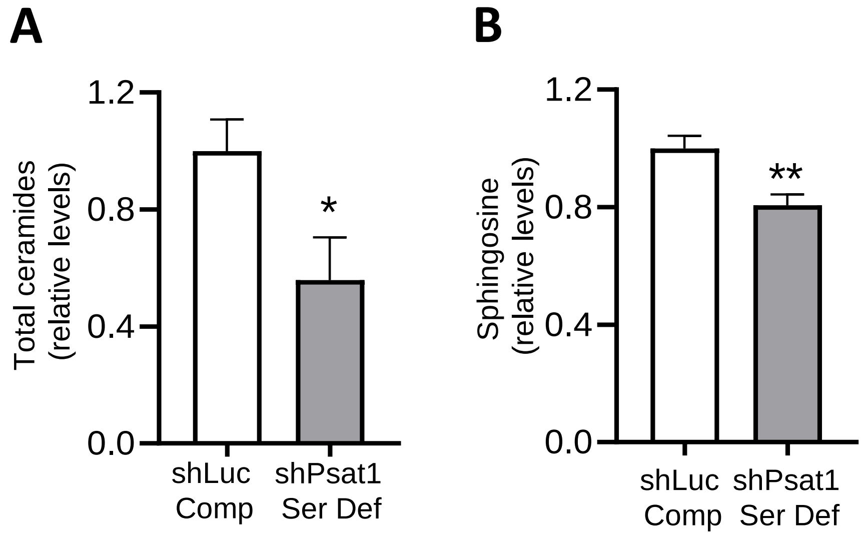


**Figure S2. Canonical sphingolipid levels do not increase following serine deprivation in colon cancer cells (related to Figure 2).**

**A-B,** CT26 cells carrying doxycycline (doxy)-inducible shRNAs targeting *Luciferase* (*shLuc*) or *Psat1* (*shPsat1)* were cultured in complete (Comp) or serine deficient (Ser Def) medium, as indicated, and treated with doxy, for 24 h. The abundance of total ceramides (**A**) and sphingosine (**B**) was determined and displayed as relative to the *shLuc* Comp group. Data represent mean values ± S.D. *P≤0.05; **P≤0.01; ***P≤0.001 (paired t-tests).

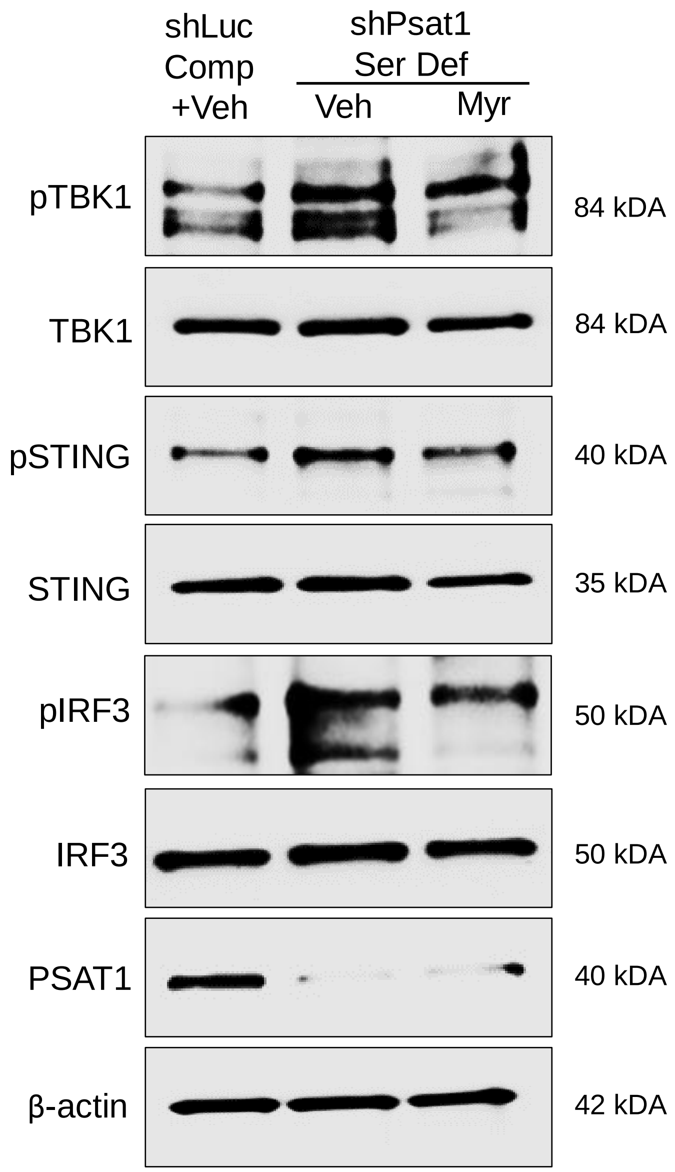


**Figure S3. Myriocin treatment attenuates the activation of cGAS-STING1 induced by serine deficiency (related to Figure 2).**

CT26 cells carrying doxycycline (doxy)-inducible shRNAs targeting *Luciferase* (*shLuc*) or *Psat1* (*shPsat1)*, were cultured in complete (Comp) or serine deficient (Ser Def) medium, respectively, and treated with vehicle (Veh) (0.1% methanol) or 500 nM myriocin (Myr), as indicated, for 24 h. Cells were harvested and probed for the indicated proteins.

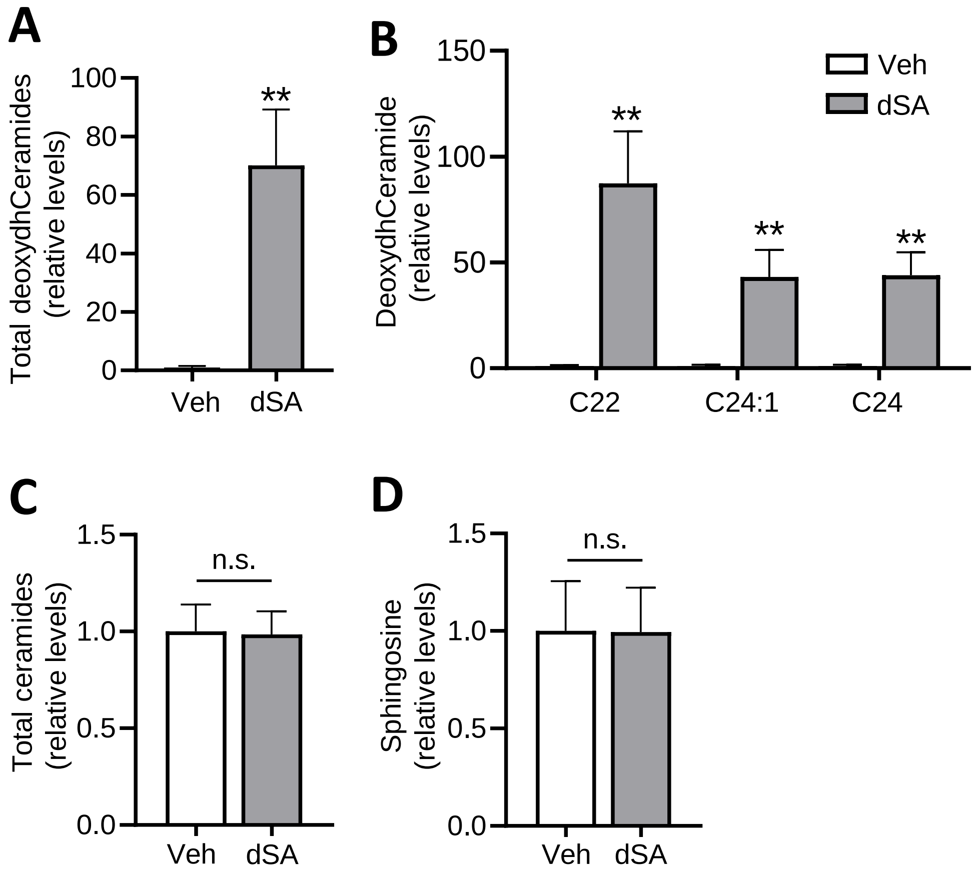


**Figure S4. Deoxysphinganine treatment increases deoxySL levels in colon cancer cells (related to Figure 3).**

**A-D,** WT CT26 cells were treated with vehicle (Veh) (DMSO) or 400 nM of 1-deoxysphinganine (dSA), for 24 h and the abundance of total deoxydihydroCeramides (deoxydhCeramides) (**A**), individual deoxydhCeramide species (**B**), total ceramides (**C**) and sphingosine (**D**) was measured and displayed as relative to the Veh group. Data represent mean values ± S.D. *P≤0.05; **P≤0.01; ***P≤0.001, (paired t-tests).

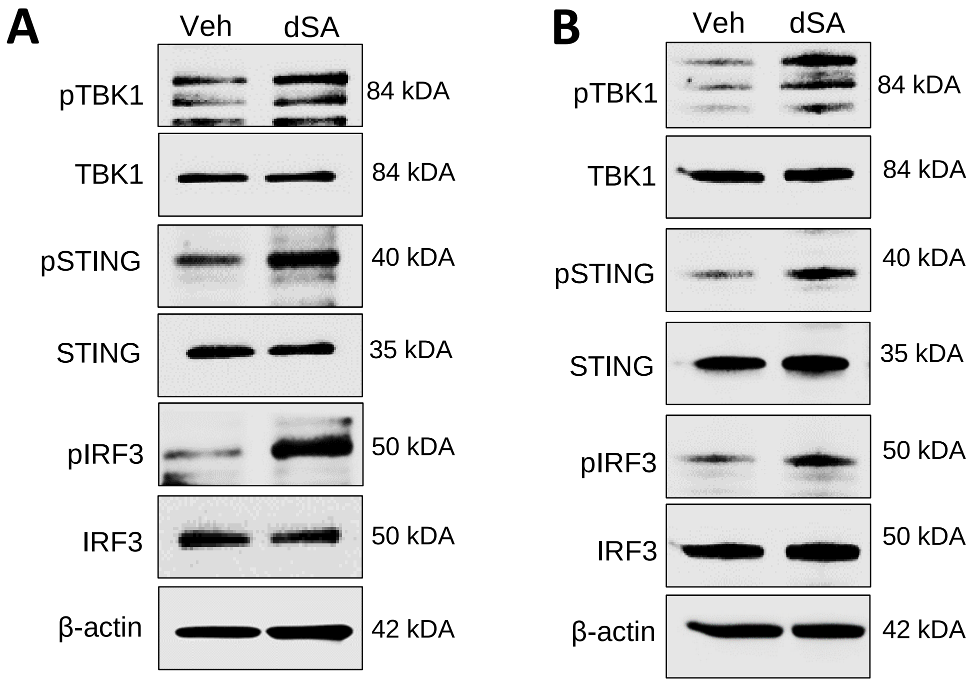


**Figure S5. Deoxysphinganine treatment induces activation of the cGAS-STING1 pathway (related to Figure 3).**

**A**, CT26 WT cells were treated with vehicle (Veh) (DMSO) or 500 nM 1-deoxysphinganine (dSA) for 18 h and subjected to western blotting analysis to probe for the indicated proteins. **B**, CT26 cells were treated and protein analyzed as described in panel A, in an independent experiment.

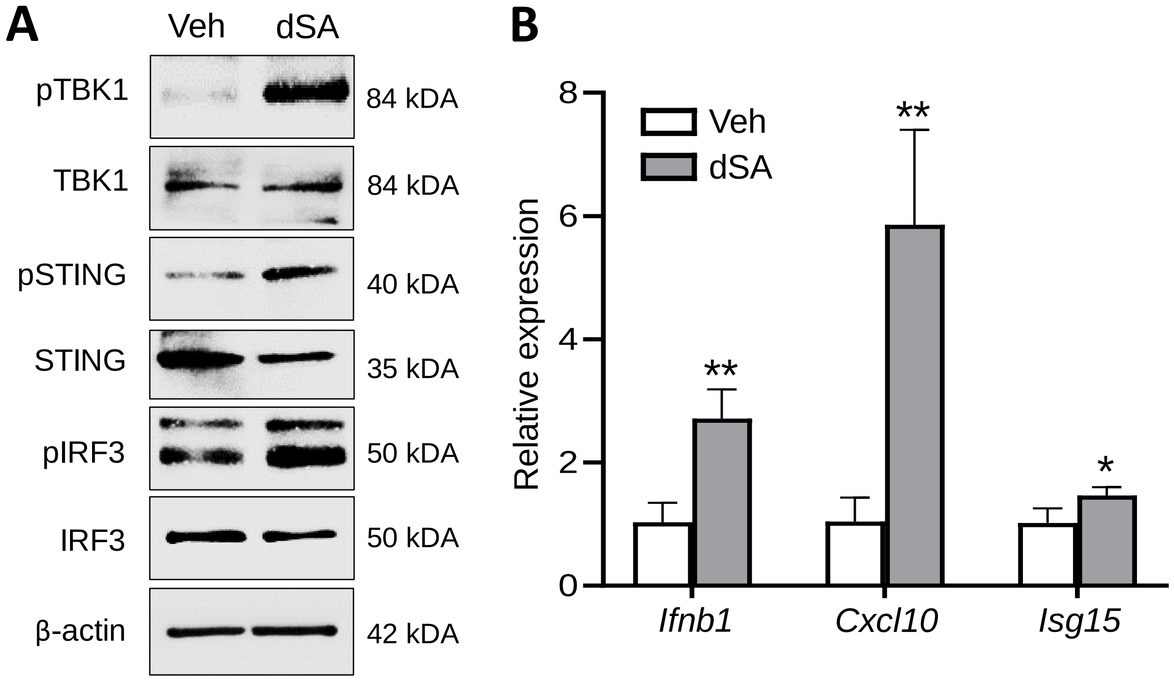


**Figure S6. Deoxysphinganine treatment triggers the induction of cGAS-STING1 in human colon cancer cells (related to Figure 3).**

**A,** DLD1 cells were treated with vehicle (Veh) (DMSO) or 500 nM 1-deoxysphinganine (dSA) for 18 h and probed for the indicated proteins. **B,** Cells were treated as described in panel a for 24 h and expression of the indicated genes was determined. Data represent mean values ± S.D. *P≤0.05; **P≤0.01 (paired t-tests).

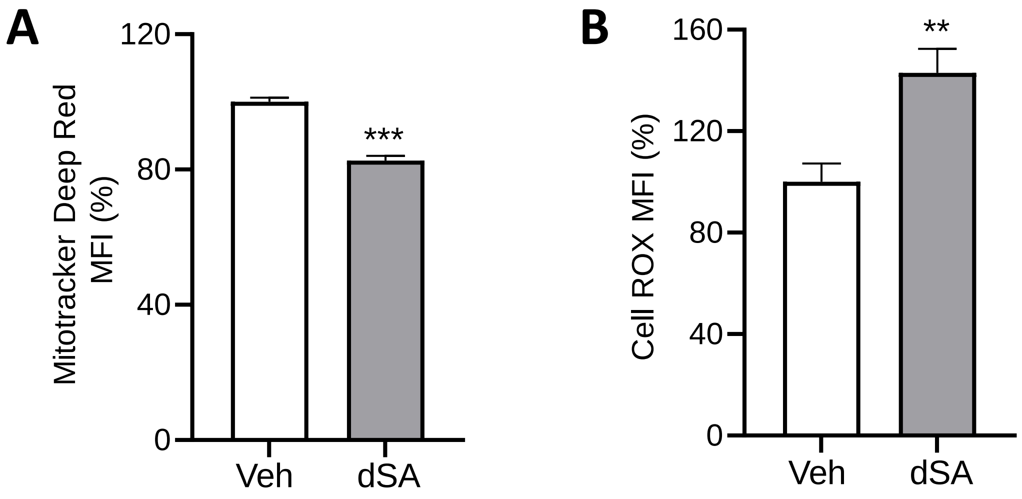


**Figure S7. Deoxysphinganine treatment promotes mitochondrial dysfunction in DLD1 cells (related to Figure 4).**

**A-B**, DLD1 cells were treated with vehicle (DMSO) or 500 nM of 1-deoxysphinganine (dSA) for 6 h and mitochondrial membrane potential was determined after staining with Mitotracker Deep Red (**A**) and reactive oxygen species levels were determined after staining with CellROX (**B**), by flow cytometry. Data represent mean values ± S.D. ***P≤0.001.

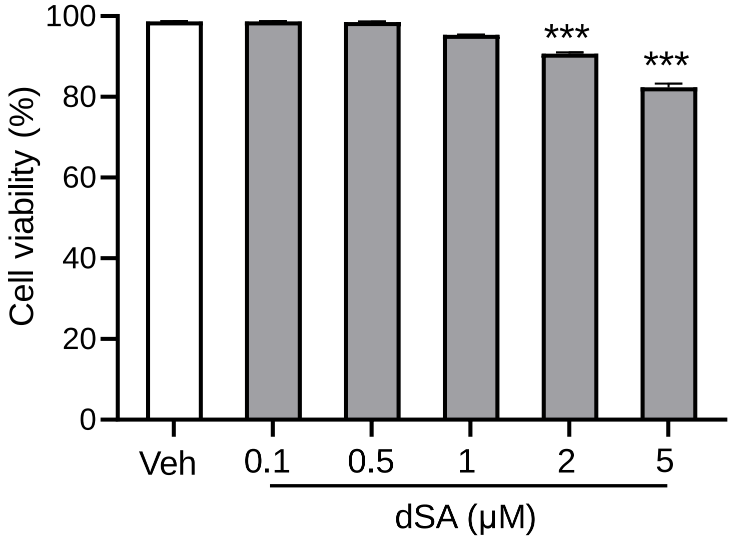


**Figure S8. Concentration of deoxysphinganine used to induce cGAS-STING1 does not reduce viability of CT26 cells (related to Figure 4).**

WT CT26 cells were treated with vehicle (Veh) (DMSO) and 1-deoxysphinganine (dSA) (0.1, 0.5, 1, 2 and 5 µM) for 24 h and viability was determined by flow cytometry following DiOC6 and DAPI staining. Data represent mean values ± S.D.

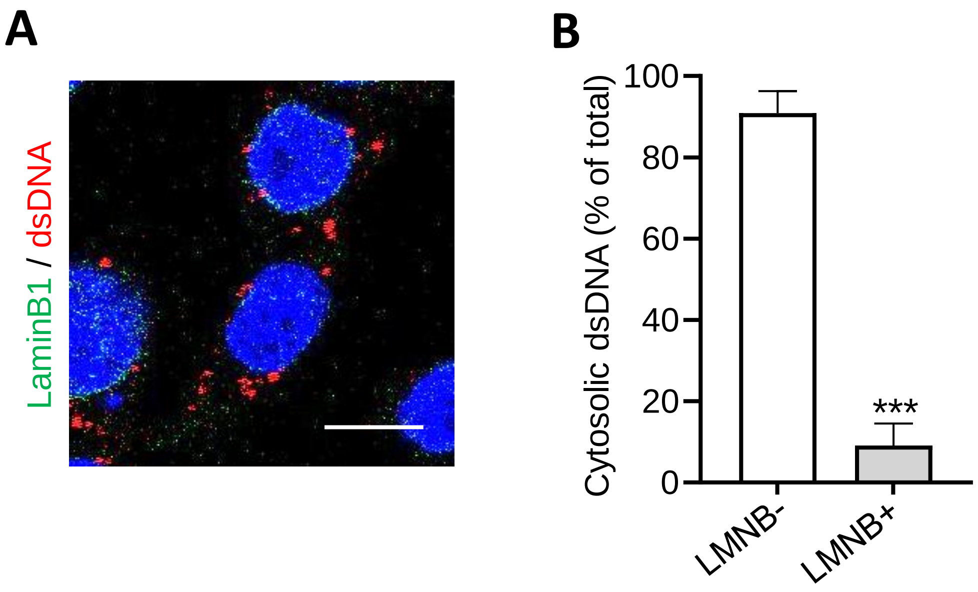


**Figure S9. Minimal cytosolic dsDNA associated with the nucleus following deoxysphinganine treatment of CT26 cells (related to Figure 4).**

**A-B,** WT CT26 cells were treated with 500 nM of 1-deoxysphinganine (dSA) for 6 h followed by staining with antibodies specific for dsDNA (red color) and Lamin B (to identify nuclear envelope) (green color). DAPI (blue color) was used to stain the nucleus. Representative images (**A**) and quantification of the percent of cytosolic dsDNA dots in proximity with Lamin B-positive structures (**B**), are shown. Scale bar = 10 μm. Data represent mean values ± S.D. ***P≤0.001, (Fisher's t-test).

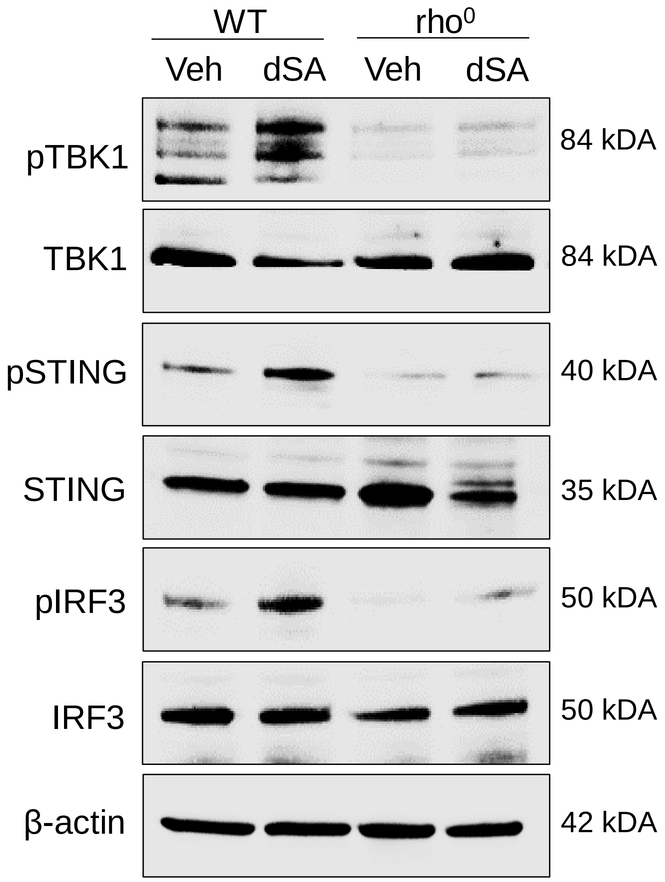


**Figure S10. Deoxysphinganine treatment fails to elicit cGAS-STING1 pathway activation in rho$^0$ cells (related to Figure 4).**

CT26 WT and rho$^0$ cells were treated with vehicle (Veh) (DMSO) or 500 nM of 1-deoxysphinganine (dSA) for 18 h and probed by western blot for the indicated proteins.

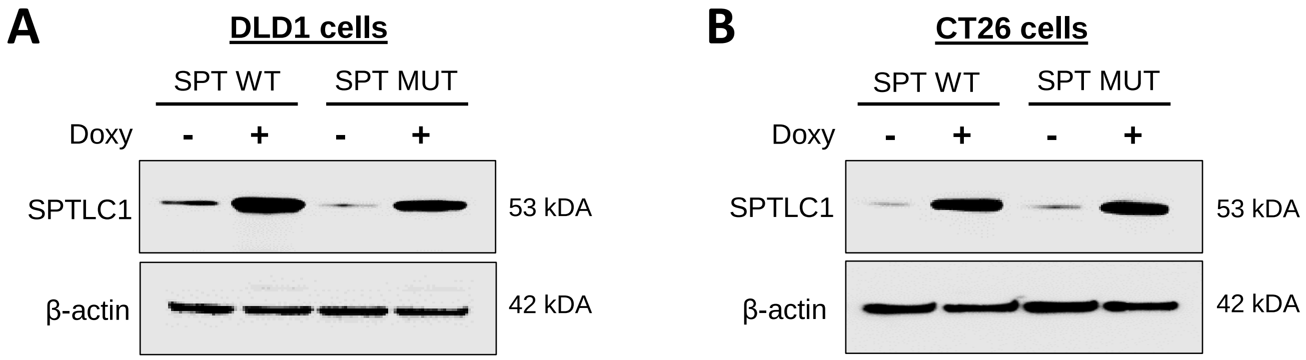


**Figure S11. Doxycycline-induced expression of WT or mutant SPT in CT26 and DLD1 cells (related to Figure 5).**

**A**, Doxycycline (doxy)-inducible *SPTLC1* wild-type (SPT WT) and *SPTLC1*$^{C133W}$ mutant (SPT MUT) DLD1 cells were treated with or without doxy for 24 h and probed for the indicated proteins.

**B**, Doxycycline (doxy)-inducible CT26 *SPTLC1* wild-type (SPT WT) and mutant (SPT MUT) cells were treated as described in panel a and probed for the indicted proteins using western blotting.

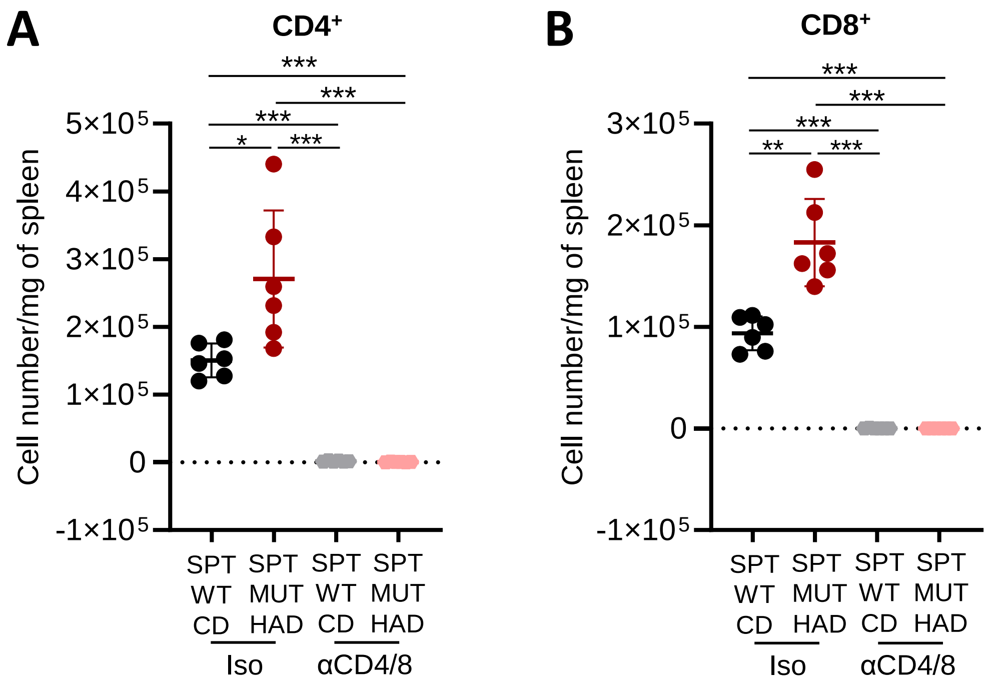


**Figure S12. Reduced immune cell abundance in spleens of mice treated with anti-CD4 and CD8 blocking antibodies (related to Figure 7)**.

**A-B,** Doxycycline (doxy)-inducible *SPTLC1* wild-type (SPT WT) and *SPTLC1*$^{C133W}$ mutant (SPT MUT) CT26 cells were injected subcutaneously into BALB/c mice and tumors were allowed to grow to a palpable size while being fed control complete diet (CD). Both groups of tumor-bearing mice were then randomized to receive either isotype control (Iso) or CD4 and CD8 blocking antibodies (α-CD4/8) on days 8 and 16. All mice received 20 mg/kg of doxy by oral gavage every other day to activate shRNAs beginning on day 10, and SPT WT tumor bearing-mice were given CD while SPT MUT tumor-bearing mice were given a high alanine diet (HAD). The total number of CD4$^{+}$ T cells **(A)** and CD8$^{+}$ T cells **(B)** per mg of spleen was determined (n=6 mice per group). Data represent mean values ± S.D. *P≤0.005; **P≤0.01; ***P≤0.001 (paired t-tests).

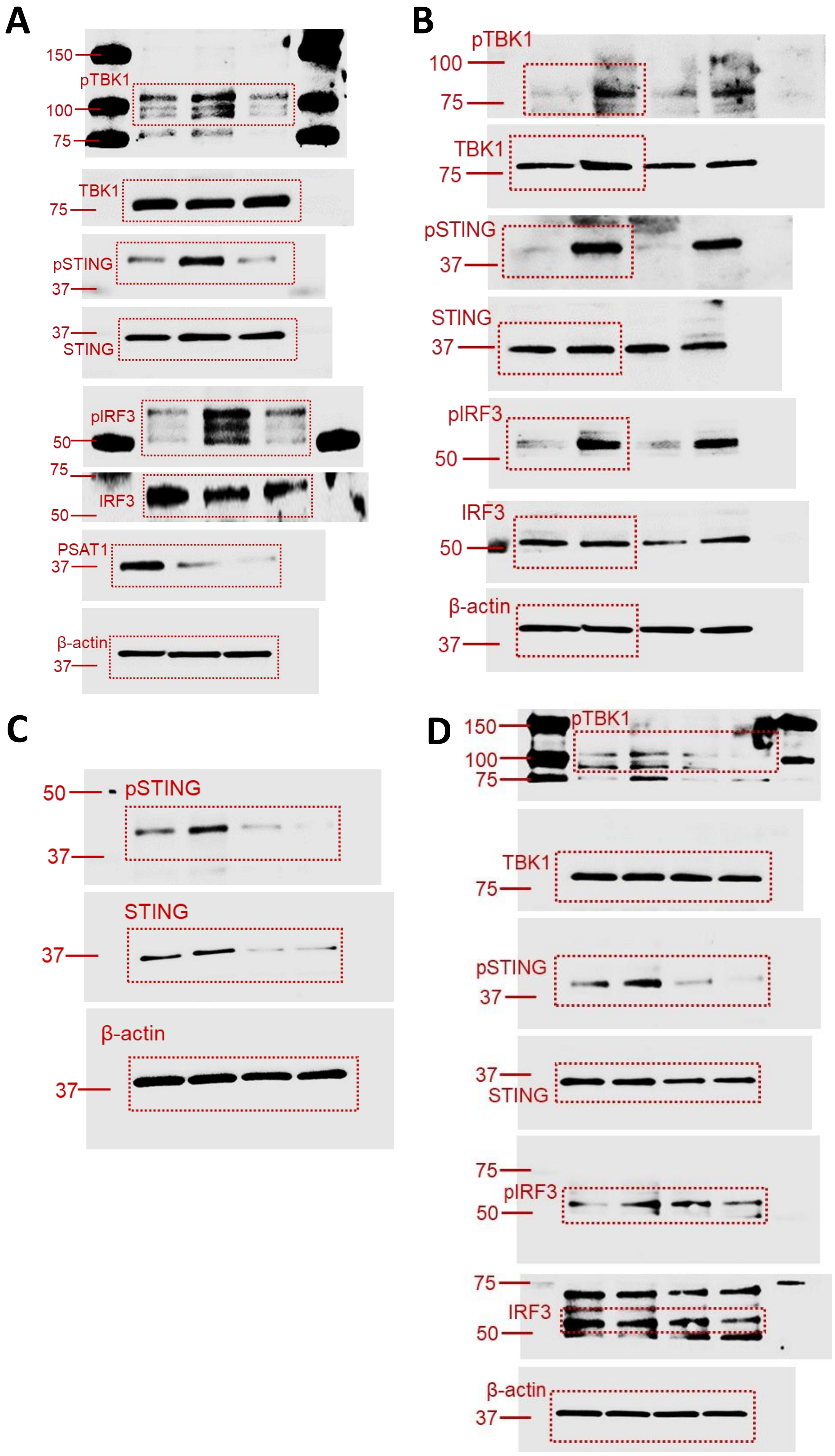


**Figure S13. Uncropped images of western blots (related to Figures 2, 3 and 4).**

**A-D,** Images of original uncropped membranes of Western blots presented in Figure 2D (**A**), Figure 3A (**B**), Figure 3E (**C**) and Figure 4F (**D**). Red boxes depict the areas of the blots shown in main figures.

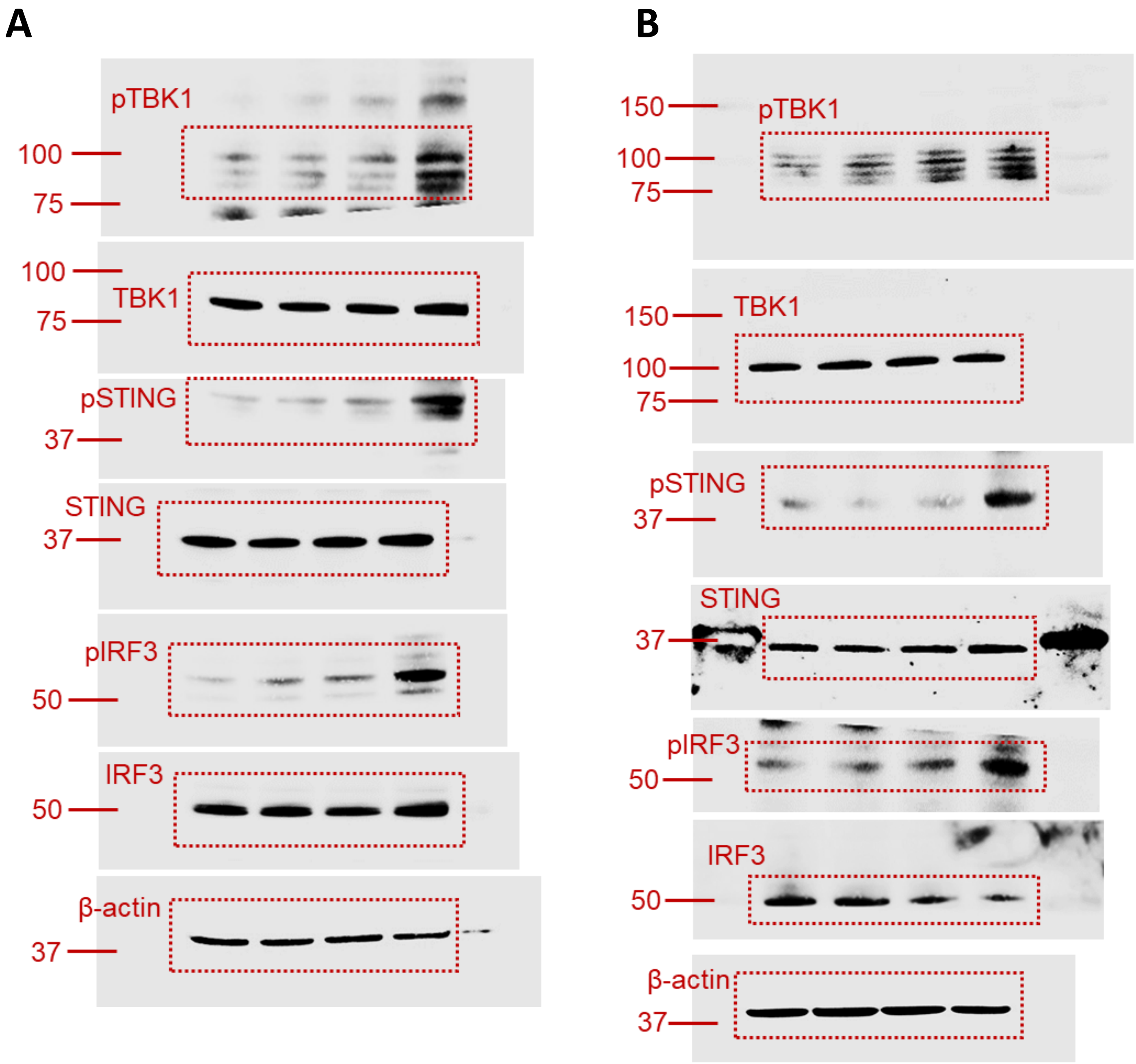


**Figure S14. Uncropped images of western blots (related to Figure 5).**

**A-B** Images of original uncropped membranes of Western blots presented in Figure 5B (left side) (**A**) and Figure 5B (right side) (**B**). Red boxes depict the areas of the blots shown in main figures.

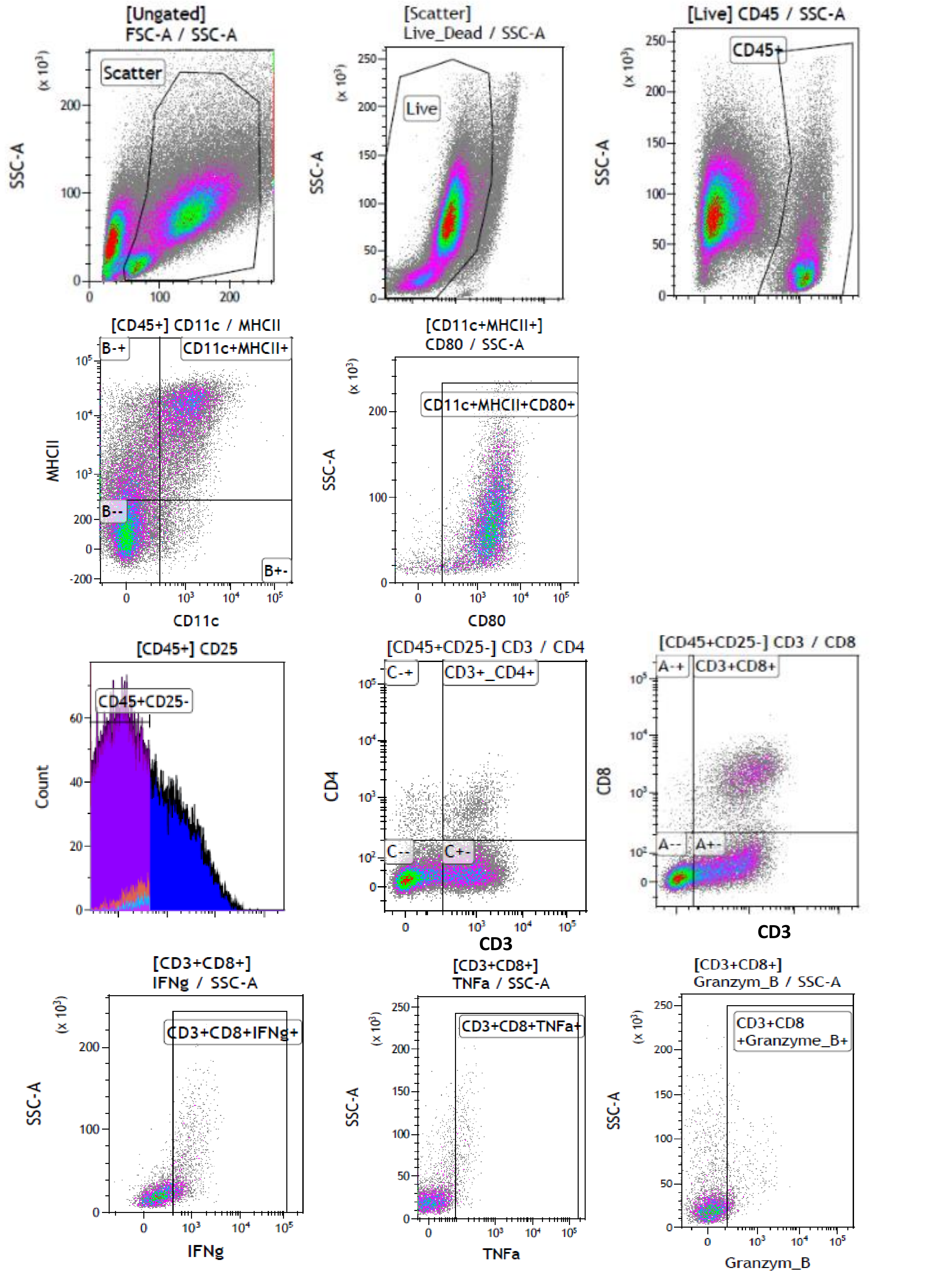


**Figure S15. Representative FACS plots for analysis of activated dendritic and T cells (related to Figures 1 and 7).** Representative plots generated by FACS analysis used to identify and quantify immune cell populations shown in Figure 1H, 7C and 7E are shown. Immune cell subsets were gated off of live CD45$^{+}$ population.

**Supplementary Table 1.** Composition of amino-acid based complete and high alanine diets.

| | Complete diet | High alanine diet |
|---|---|---|
| | gm% | gm% |
| **Ingredient** | gm | gm |
| L-Arginine | 6 | **4.5** |
| L-Histidine-HCl-H2O | 4.6 | 4.6 |
| L-Isoleucine | 7.6 | 7.6 |
| L-Leucine | 15.8 | 15.8 |
| L-Lysine-HCl | 13.2 | 13.2 |
| L-Methionine | 5.1 | 5.1 |
| L-Phenylalanine | 8.4 | 8.4 |
| L-Threonine | 7.2 | 7.2 |
| L-Tryptophan | 2.1 | 2.1 |
| L-Valine | 9.3 | 9.3 |
| L-Alanine | 5.1 | **30** |
| L-Asparagine-H2O | 6.7 | **5** |
| L-Aspartate | 5.4 | **4.1** |
| L-Cystine | 4.2 | **3.2** |
| L-Glutamic Acid | 21.7 | **16.3** |
| L-Glutamine | 16.5 | **12.4** |
| Glycine | 3 | **2.3** |
| L-Proline | 17.8 | **13.4** |
| L-Serine | 10 | **7.5** |
| L-Tyrosine | 9.2 | **6.9** |
| **Total L-Amino Acids** | **178.9** | **178.9** |
| Casein | 0 | 0 |
| Corn Starch | 397.486 | 397.486 |
| Maltodextrin 10 | 132 | 132 |
| Sucrose | 107.0777 | 107.0777 |
| Cellulose | 50 | 50 |
| Soybean Oil | 70 | 70 |
| t-butylhydroquinone | 0.014 | 0.014 |
| Mineral Mix S10022C | 3.5 | 3.5 |
| Calcium Carbonate | 7.34 | 7.34 |
| Potassium Citrate, 1 H20 | 2.4773 | 2.4773 |
| Potassium Phosphate, Monobasic | 6.86 | 6.86 |
| Calcium Phosphate, dibasic | 7 | 7 |
| Sodium Chloride | 2.59 | 2.59 |
| Sodium Bicarbonate | 7.5 | 7.5 |
| Vitamin Mix V10037 | 10 | 10 |
| Choline Bitrartrate | 2.5 | 2.5 |
| **Total** | **985.245** | **985.245** |
| Protein (gm) | 179 | 179 |
| Carbohydrate (gm) | 647 | 647 |
| Fat (gm) | 70 | 70 |
| Fiber (gm) | 50 | 50 |
| Protein (kcal) | 716 | 716 |
| Carbohydrate (kcal) | 2586 | 2586 |
| Fat (kcal) | 630 | 630 |
| **Total** | **3932** | **3932** |
| Protein (gm%) | 18 | 18 |
| Carbohydrate (gm%) | 66 | 66 |
| Fat (gm%) | 7 | 7 |
| Protein (kcal%) | 18 | 18 |
| Carbohydrate (kcal%) | 66 | 66 |
| **Fat (kcal%)** | **16** | **16** |
| **Kcal/g** | **3.99** | **3.99** |
